# Rejuvenation rankings from CRISPRi screens change with the aging cohort and track cell growth programs

**DOI:** 10.64898/2026.11.11.766060

**Authors:** Md Kawsar, Sajib Acharjee Dip, Bikash Kumar Paul, Abdullah Al Noman, Liqing Zhang, Mohammad Ali Moni

## Abstract

Ranking gene knockdowns by how well they reverse an aging expression signature is a common way to find rejuvenation targets, but the stability of these rankings has not been tested. We built TRACE, an open-source set of tests, to score 18,746 knockdown profiles from five CRISPR interference (CRISPRi) screens in four human cell lines. Rankings built from fibroblasts of 111 donors and from Genotype-Tissue Expression (GTEx) fibroblasts were unrelated (ρ = −0.05). In an exploratory test, the 200 strongest age genes of the first cohort changed in the opposite direction in GTEx (ρ = −0.55), but this was not significant after correction and shrank when GTEx batch was modeled. Raw scores followed the broad programs inside each signature: strongly for proliferation with the fibroblast signature, weakly for stress with a GTEx-derived signature. Proliferation alone matched the fibroblast scores’ agreement with senescence labels. TRACE runs on other aging cohorts and screens.

**MOTIVATION:** Perturbation atlases make it easy to rank thousands of genes by how well their knockdown reverses an aging signature. But there is no standard way to check whether such a ranking depends on the cohort behind the signature or simply follows broad programs such as proliferation. TRACE swaps the aging cohort, checks whether the cohort test could detect a shared effect, compares score definitions with random signatures and a proliferation-only baseline, and tests rankings against held-out labels.

## INTRODUCTION

Aging is accompanied by coordinated changes in gene expression that reflect genomic instability, loss of proteostasis, mitochondrial dysfunction, cellular senescence and altered communication between cells^1^. These changes can be measured cheaply and at scale, so transcriptomic signatures of age have become a common read-out for interventions that might slow or reverse aging^2^. In this setting, rejuvenation means a shift of a cell’s molecular state toward that of a younger cell, and it is usually judged by such read-outs rather than by function^3,4^. Because experiments on aging are slow and costly, rankings built on these read-outs decide which candidates are tested.

This strategy builds on the connectivity map, in which a disease or aging signature lists the genes that go up and down with the condition, and perturbations whose expression response runs opposite to that signature are ranked highest^5,6^. The approach has nominated geroprotective compounds such as HSP90 inhibitors^7^. Related signatures of lifespan-extending interventions have been compared across tissues and species^8,9^. Newer work scales the same logic to large perturbation libraries. REVIVE combines a transcriptional aging clock with profiles of more than 10,000 chemical and genetic perturbations^10^, and a fine-tuned single-cell foundation model has been used to nominate anti-aging gene knockdowns in silico^11^. Interest has grown further because partial reprogramming can reverse molecular and functional features of aging in vivo^12,13^. These approaches assume that the ranking reflects aging biology rather than the particular data used to build it.

Several observations suggest that this assumption may not always hold. Connectivity Map signatures reproduce poorly between versions of the resource^14^. Genes that change with age also overlap little across tissues and datasets^15^. CRISPR interference (CRISPRi)^16^ Perturb-seq screens now measure transcriptome-wide responses to the knockdown of thousands of genes in several human cell lines^17,18^. Models trained on these screens, such as GEARS^19^ and scGPT^20^, are proposed as engines for in silico target discovery. As a result, reversal rankings can now be built for thousands of genes at once, but their reliability has not been tested systematically.

The reliability of such rankings depends on three issues that have not been examined together. First, an aging signature may not carry over between donor cohorts. Age effects in cultured cells can be confounded by donor source, culture history and batch^21–23^, so a signature from one cohort may partly describe that cohort. Second, part of a reversal score may simply reflect cell growth. Aged and senescent cells divide less^24,25^, and aging transcriptomes across human tissues track reduced proliferation^26^. Although proliferation is part of aging biology, it is also a strong confounder of expression signatures, to the point that most random signatures predict breast cancer outcome through it^27^. A knockdown that speeds growth may look young, although growth-promoting changes can be oncogenic^28^. Third, it is unclear whether a ranking agrees across cell lines and with independent evidence better than simple baselines do. In perturbation modeling, simple baselines are often hard to beat^29^. Earlier rejuvenation screens aimed to nominate candidates^7,10,11^. None, to our knowledge, swapped the aging cohort, compared each score with a proliferation-only baseline, or asked whether a cohort test could detect agreement if it existed.

Here we introduce TRACE (Transferable Rejuvenation Assessment across Contexts and Evidence). Rather than proposing a new ranking method, TRACE provides a set of tests that any reversal ranking can be run through from a single command (Figure 1A). It rebuilds the ranking with a signature from a second cohort and checks with a planted effect whether this cohort test has power. It compares several score definitions, including ones adjusted for proliferation, against random signatures and a proliferation-only baseline. It measures agreement between cell lines against agreement between replicate screens. Finally, it tests rankings against held-out senescence and longevity labels. We fixed five hypotheses and their tests before scoring any knockdown (Table 1), and we label all later analyses as exploratory. We applied TRACE to a widely used fibroblast aging signature^21^, a second signature built the same way from GTEx fibroblasts^22^, and five CRISPRi screens in four human cell lines^17,18^.

**Figure 1.**
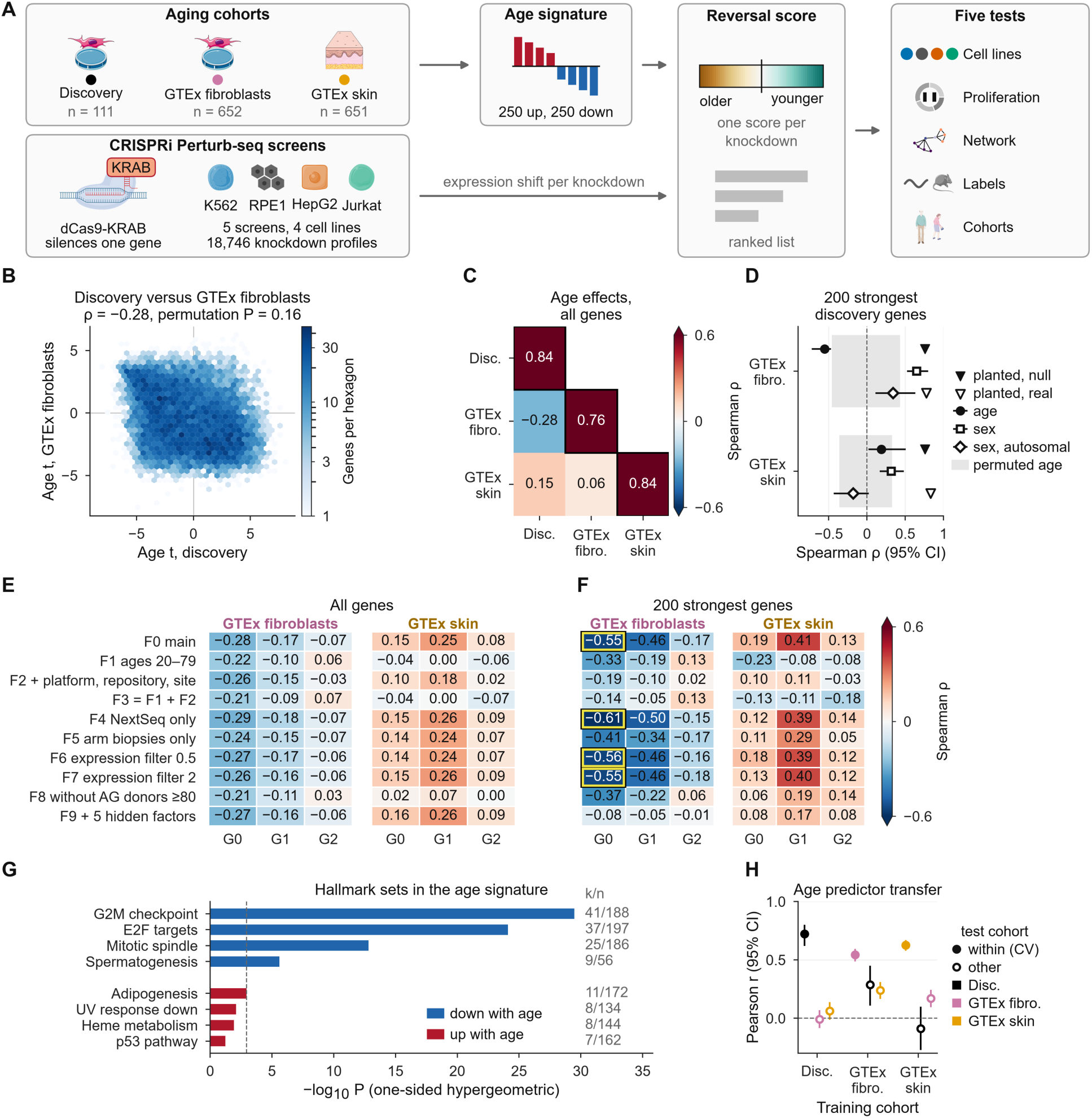
Study design and cross-cohort transfer of the fibroblast age signature. (A) TRACE workflow: cohorts and screens give an age signature and a reversal score per knockdown, which five tests check. (B) Age t-statistics in the discovery cohort versus GTEx cultured fibroblasts (10,538 shared genes); ρ, Spearman correlation; P, two-sided permutation P. (C) Agreement of age effects (Spearman ρ) between cohorts (off-diagonal) and between split halves of one cohort (diagonal, outlined; median); color scale as in (F). (D) ρ of age and sex t-statistics for the 200 strongest discovery genes (sex: all and autosomal genes); error bars, 95% bootstrap CIs; gray band, 95% range under permuted age; triangles, median ρ for a shared effect planted after permuting age (filled) or added to the real data (open). (E and F) ρ over all genes (E) and the 200 strongest discovery genes (F) for each discovery specification (rows) against GTEx fibroblasts and skin under models G0 to G2 (columns); black-and-yellow outline, G0 cells outside their own permuted-age null. AG, NIA Aging Cell Repository. (G) Hallmark gene sets most enriched among down- and up-regulated signature genes (one-sided hypergeometric test; k/n, overlap/set size; dashed line, FDR = 0.05). (H) Pearson r between elastic-net predicted and true age within each training cohort (cross-validated, filled) and in other cohorts (open; color, test cohort); error bars, 95% CIs. The P in (B) and panels D to F and H are exploratory. See also Figures S1 and S2 and Tables S2 to S8.

**Table 1.** Hypotheses fixed before analysis and their outcomes.

| ID | Hypothesis | Test and decision rule | Holm family | Result | Outcome |
|---|---|---|---|---|---|
| H1a | Raw scores correlate positively between every pair of lines | One-sided Spearman per pair; all pairs significant | H1a (6 pairs) | ρ = 0.42–0.64 | Supported |
| H1b | Within-line (two K562 screens) agreement exceeds between-line agreement | Bootstrap (2,000); one-sided 95% bound > 0 | – | 0.50 vs 0.51 | Not supported |
| H2 | Raw scores correlate with proliferation shift in every screen | Spearman per screen | H2 (5 screens) | ρ = 0.49–0.90 | Supported |
| H3a | Network beats degree-preserving randomized networks | Paired one-sided Wilcoxon on absolute error; majority of screens | H3a (5 screens) | 3/5 screens (RPE1, HepG2, Jurkat) | Supported<sup>a</sup> |
| H3b | Line transfer + network beats uncalibrated line transfer | As H3a | H3b (5 screens) | 5/5 screens | Supported<sup>b</sup> |
| H4a | CellAge inducers score higher than inhibitors | One-sided Mann–Whitney on R<sub>rob</sub> | H4 (2 tests) | AUROC 0.54 | Not supported |
| H4b | GenAge anti-longevity orthologs score higher than pro-longevity orthologs | As H4a | H4 (2 tests) | AUROC 0.51 | Not supported<sup>c</sup> |
| H5a | Donor-bootstrap stability of R<sub>rob</sub> | Median ρ over 200 bootstraps ≥ 0.8 | – | Median ρ = 0.779 | Not supported<sup>d</sup> |
| H5b | Ranking from the GTEx-derived signature correlates positively with the discovery ranking | One-sided Spearman, α = 0.05 | – | ρ = −0.05 | Not supported |
| H5 | Overall (H5a and H5b both required) | – | – | – | Not supported |

Holm families were not combined across hypotheses. AUROC, area under the receiver operating characteristic curve.
<sup>a</sup>Exploratory checks: against the mean of 500 randomizations the error reduction was significant in 2 of 5 screens; the bootstrap CI of the mean error difference excluded zero only in HepG2; in K562 genome-wide the network had 6.3% higher error than the null.
<sup>b</sup>The reference in the plan is uncalibrated line transfer; most of the gain comes from recalibration (exploratory comparison with calibrated line transfer in the Results).
<sup>c</sup>A post hoc fix to the H4b input (an ortholog table skipped by a file-size check) changed the AUROC from 0.52 to 0.51 (Table S19).
<sup>d</sup>Exploratory check: 2,000 bootstraps gave a median of 0.778.

We found that a reversal ranking depends on the cohort used to build its signature. Rankings built from the two fibroblast cohorts were unrelated, and this held for every score definition. Two random halves of a single cohort agreed more closely than the two cohorts did. Raw scores also followed the broad expression programs contained in each signature. The fibroblast signature contains many cell-cycle genes, so its scores tracked how strongly each knockdown slowed growth. The GTEx signature contains few, and its scores did not track growth in the same way. For both signatures, agreement with senescence labels was no better than proliferation alone. A small signal shared across cell lines remained after proliferation was removed, indicating that the scores are not pure noise. Based on these results, we propose a short checklist for any transcriptomic rejuvenation screen (Box 1): build the ranking from a second cohort, show that the cohort test has power, and report every candidate against a proliferation-only baseline before calling it a rejuvenation target.

## RESULTS

### The strongest fibroblast age genes do not transfer between cohorts

We assembled three aging cohorts and five CRISPRi screens (Figures 1A and S1A; Table S2). The discovery cohort comprised primary dermal fibroblasts from 111 healthy adult donors^21^. The replication cohorts were cultured fibroblasts (652 donors) and non-sun-exposed skin (651 donors) from release v10 of GTEx^22,30^. GTEx ages are available only in 10-year bins from 20 to 79 years. By contrast, 33 discovery donors were aged 80 or older, and 31 of them came from one repository (AG, the Aging Cell Repository of the National Institute on Aging, NIA), so age and repository overlap (Table S3; STAR Methods).

Of 11,732 expressed genes, 6,495 were associated with age in the discovery cohort (expression ∼ age + sex; Benjamini–Hochberg false discovery rate, FDR < 0.05)^31^, against a median of 0 under permuted age labelings (Figure S1B). Limiting the cohort to donors aged 20 to 79 left 198 genes, against a median of 5,128 in random subsets of the same size (Figure S1C), and removing the 31 AG donors aged 80 or older left none (exploratory). The signal comes from the oldest donors, most of them from one repository, and the two cannot be separated. The signature comprised the 250 most up-and 250 most down-regulated genes.

Genome-wide, the Spearman correlation of age t-statistics between the discovery cohort and GTEx fibroblasts was ρ = −0.28 (Figure 1B), and with GTEx skin it was 0.15 (Figure 1C). Neither differed from chance (two-sided permutation *P* = 0.16 and 0.28; Figure S1D). Exploratory P values in the text are uncorrected unless a q value is given. But this test has little power: thousands of genes share a few expression programs, so even permuted ages give correlations as large as ±0.4. GTEx fibroblasts and skin also agreed poorly (ρ = 0.06; *P* = 0.57), whereas two random halves of one cohort agreed well (Figure 1C), an upper bound because halves share platform, repository and processing.

We therefore also compared the 200 strongest discovery age genes, an exploratory test chosen after the first results because 200 is close to the size of each signature arm. Their correlation with GTEx fibroblasts was −0.55 (permutation *P* = 0.011; Figure 1D), and ρ stayed −0.57 to −0.50 for the 50 to 1,000 strongest genes, outside the permuted-age 95% range at every k (Figure S1E); with GTEx skin it stayed inside that range. The reversal fell inside the null in six specifications, mostly those without the oldest, single-repository donors, which are also smaller, but it never turned positive under the main GTEx model (G0; Table S4; STAR Methods). The original model (F0) is the main exploratory comparison; the others are sensitivity checks. Across all 47 cohort tests no P value survives correction (q = 0.17 for F0; Table S5). Cell-cycle genes explain part of the reversal: Hallmark^32^ E2F-target and G2M-checkpoint genes and *MKI67*, *TOP2A* and *LMNB1* fell with age in the discovery cohort but rose in GTEx fibroblasts (Figures S1F, S1G and S2; Table S6).

As a positive control, sex effects agreed among the 200 most sex-biased discovery genes (ρ = 0.65 with GTEx fibroblasts, permuted-sex *P* = 0.002; 0.32 with skin, *P* = 0.057), and every canonical sex gene had the same sign in all three cohorts, which rules out a merge or sign error (Table S6). But this agreement came from sex-chromosome genes (Figure 1D; Table S7), so it cannot show that the pipeline would recover a moderate autosomal effect such as aging. We therefore planted a shared effect in 500 genes of both cohorts after permuting age (median discovery |t| = 2.1 and 4.3; 200 replicates each). The 200 strongest discovery genes recovered it in every replicate (median ρ = 0.73–0.76; Figure 1D), but the genome-wide correlation exceeded its null in only 5–11% of replicates, so the genome-wide test cannot show that a shared age effect is absent. Added to the real data, the shared effect gave a positive top-200 ρ in 100% of runs (Table S3). So a shared effect of this size would have been seen, but a weaker or sparser one, or one that appears only after age 80 (GTEx has no donors that old), could be hidden.

Crossing ten discovery specifications (F0 to F9) with four replication specifications per GTEx tissue (G0 to G3) gave 80 combinations (exploratory; Figures 1E, 1F and S2; Table S4; STAR Methods). Under G0, no discovery specification agreed with either tissue genome-wide beyond its own permuted-age null (all *P* ≥ 0.16), and none gave a positive top-200 correlation with GTEx fibroblasts (ρ = −0.61 to −0.08; Figure 1F). Adding GTEx nucleic-acid isolation batch (G2) shrank the F0 reversal from −0.55 to −0.17, and 4 of 10 specifications turned positive. But G2 also removed most GTEx age genes (FDR < 0.05: 5,548 to 515 in fibroblasts, 9,778 to 31 in skin), its F0 interval was wide (−0.54 to 0.30), and with age permuted within batch the G2 value lay inside its own null (*P* = 0.40, uncorrected). Batch and age may overlap in GTEx as well. So under G2 the top-200 test can show neither a reversal nor agreement, and we cannot tell whether batch explains the reversal or hides part of a real one. Under G1, top-200 agreement with GTEx skin rose from 0.19 to 0.41 (Figure 1F). Hidden expression factors did not explain the reversal (Table S3; STAR Methods).

In short, the age signature did not transfer across the cohorts and sample types examined. Cohort composition, culture history and technical differences may all contribute, and the available metadata cannot separate them. The down-regulated arm was dominated by cell-cycle programs (Figure 1G), the part that GTEx reverses. Age-prediction models transferred no better (elastic net^33^; Figure 1H; Table S8).

### Raw reversal scores follow the programs inside the signature

For a knockdown with pseudobulk z-shift d, the raw score R is the negative Spearman correlation of d with the age t-statistics a over the signature genes S, excluding the knockdown’s own target (equation 1 in STAR Methods). R > 0 means that the knockdown moves expression opposite to the age direction (Figure 2A). We call R a transcriptomic reversal score and use “rejuvenation” only in this proxy sense. The 18,746 knockdown profiles from the K562 essential-gene (K562 ess.), K562 genome-wide (K562 gw), RPE1, HepG2 and Jurkat screens^17,18^ cover 9,870 target genes (Figure S3). We use “knockdown” for a profile in one screen and “gene” for a target pooled across screens.

**Figure 2.**
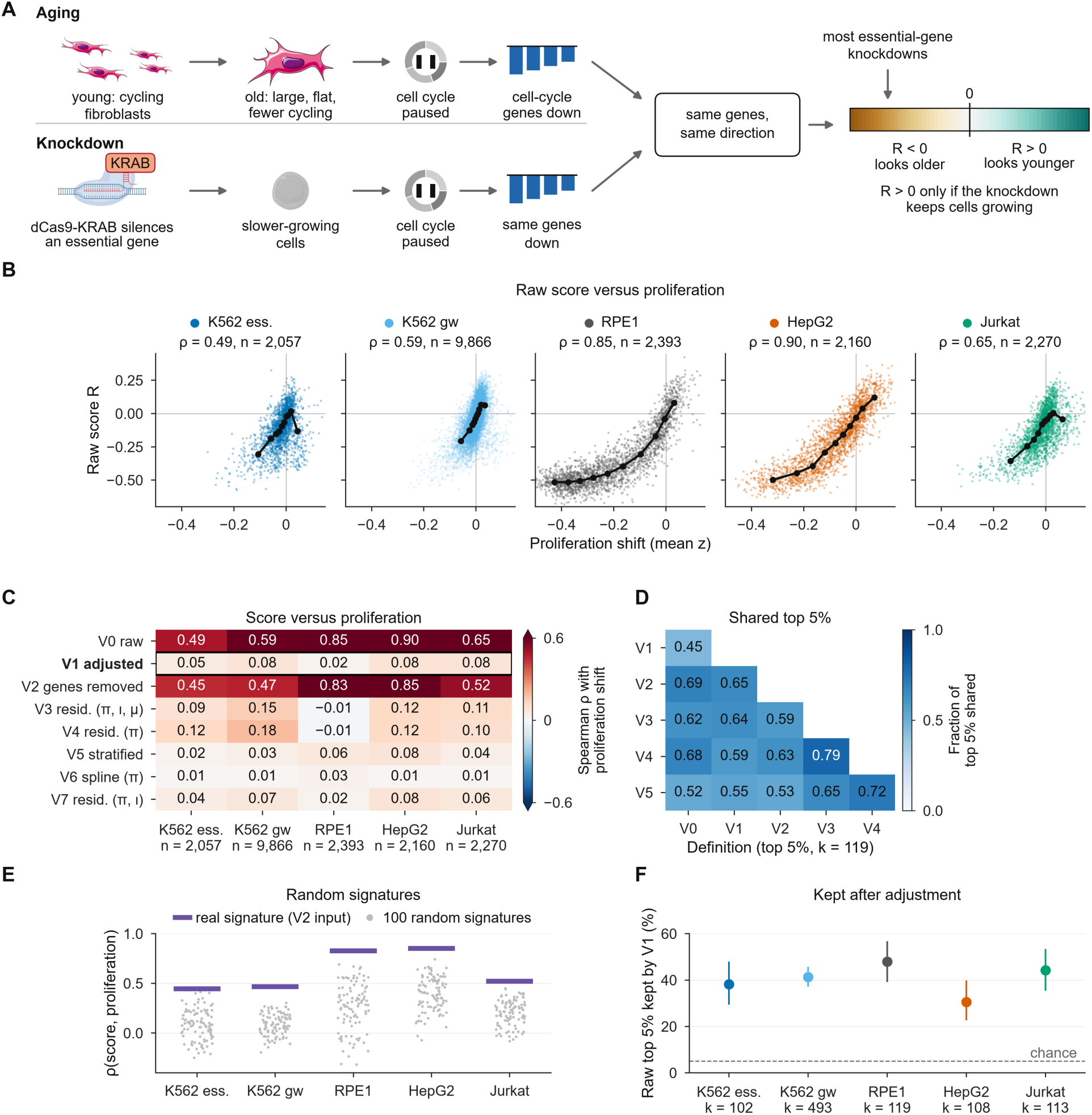
Raw reversal scores of the discovery signature track proliferation. (A) Working model: aged discovery fibroblasts express fewer cell-cycle genes, and most essential-gene knockdowns lower the same genes, so they score as “older”. (B) Raw score R versus proliferation shift, one point per knockdown; black line, decile medians; ρ and n per screen above each plot; all Holm *P* < 10^−100^. (C) Spearman ρ of each score definition with proliferation shift per screen (n, knockdowns). V0, raw; V1, main adjusted score (boxed); V2 to V7, other definitions (π, proliferation; ι, identity; μ, response size; STAR Methods). (D) Fraction of the top 5% (k = 119 of the 2,393 multi-line genes) shared between definitions V0 to V5, using R_rob_. (E) ρ with proliferation for the real signature without its proliferation genes (purple bars) and for 100 matched random signatures (dots). (F) Fraction of each screen’s raw top-5% knockdowns that stay in the top 5% under V1; error bars, binomial 95% CIs; dashed line, chance (5%). Panel B and V0 and V1 in (C) test hypotheses in Table 1; other panels are exploratory. K562 ess., K562 essential-gene screen; K562 gw, K562 genome-wide screen. See also Figures S3 to S7 and Tables S9, S10 and S11.

R correlated with each knockdown’s proliferation shift (mean z over Hallmark E2F-target and G2M-checkpoint genes) in every screen (ρ = 0.49–0.90, all Holm *P* < 10^−100^; Table 1; Figure 2B), partly through shared measurement noise (STAR Methods). R also correlated negatively with a stress and inflammation program (ρ = −0.85 to −0.47; Figure S4). In a working model (Figure 2A), most essential-gene knockdowns slow growth, aged fibroblasts in this cohort express fewer cell-cycle genes, and so a knockdown looks “aged” unless it preserves proliferation. The model rests on cell-cycle gene expression, not measured growth, and matches earlier reports that aging transcriptomes track reduced proliferation^26^.

Because proliferation is itself part of aging biology^24,25^, removing it is a choice, not a correction. We therefore compared the main adjusted score (V1) with the raw score (V0) and six exploratory definitions (V2 to V7; Figures 2C and 2D; Table S9; STAR Methods). V1 removes proliferation genes and residualizes the score on proliferation, identity-program and response-size shifts, as growth-rate corrections do in drug screens^34^. R_adj_ is the V1 value in one screen. To pool lines, the robust score R_rob_ is the mean R_adj_ over lines divided by one plus its standard deviation (SD) over lines (equation 2 in STAR Methods).

Removing proliferation genes alone left the association intact (V2: ρ = 0.45–0.85), because other signature genes co-vary with proliferation, as seen before in cancer signatures^27^. Residualization or stratification removed its monotone part (V1, V3 to V5: ρ = −0.01 to 0.18; Figure 2C), although some non-linear dependence remained for V1 in RPE1 (distance correlation^35^ 0.25; Figure S5). V1, V3, V4 and V5 shared 55–79% of their top-5% genes (Figure 2D). Table S10 lists candidates that left or stayed in the raw top 5%; none has been tested experimentally.

Is this coupling special to the aging signature? Without its proliferation genes (V2), the real signature tracked proliferation much more than 100 matched random signatures from genes outside the proliferation sets in all five screens (Figures 2E and S6; STAR Methods). It also gave higher adjusted cross-line agreement than all 100 (q = 0.014) and a higher adjusted CellAge area under the receiver operating characteristic curve (AUROC) than 97% of them (q = 0.048). So the coupling comes from the genes in this signature. A signature derived the same way from GTEx fibroblasts had almost no cell-cycle genes (2.6%, against 12.4%) and was enriched instead for oxidative phosphorylation, complement and hypoxia genes (up) and inflammatory programs (down) (Figure S7; Table S11). Its raw scores tracked stress and response size weakly and proliferation inversely. So the raw score follows the programs in its signature. For the discovery signature, 31–48% of each screen’s raw top-5% knockdowns stayed in the top 5% under V1 (Figure 2F).

### Cross-line agreement reflects the shared proliferation component

We use “line” for a cell line and “screen” for an experiment; the two K562 screens count as one line. Raw scores correlated between every pair of lines (ρ = 0.42–0.64; one-sided Holm *P* < 10^−10^; Figure 3A; Table 1), and the two K562 screens agreed at ρ = 0.49 (*n* = 2,053). On the 1,784 knockdowns measured in every screen, this within-line agreement (0.50) did not exceed between-line agreement (0.51; Table 1). After adjustment, between-line agreement fell to ρ = 0.15–0.26, below the K562 within-line value (0.38; Figure 3A), for every proliferation-adjusted definition (Figure 3B). So the raw scores’ cross-line agreement mostly reflects the shared proliferation component.

**Figure 3.**
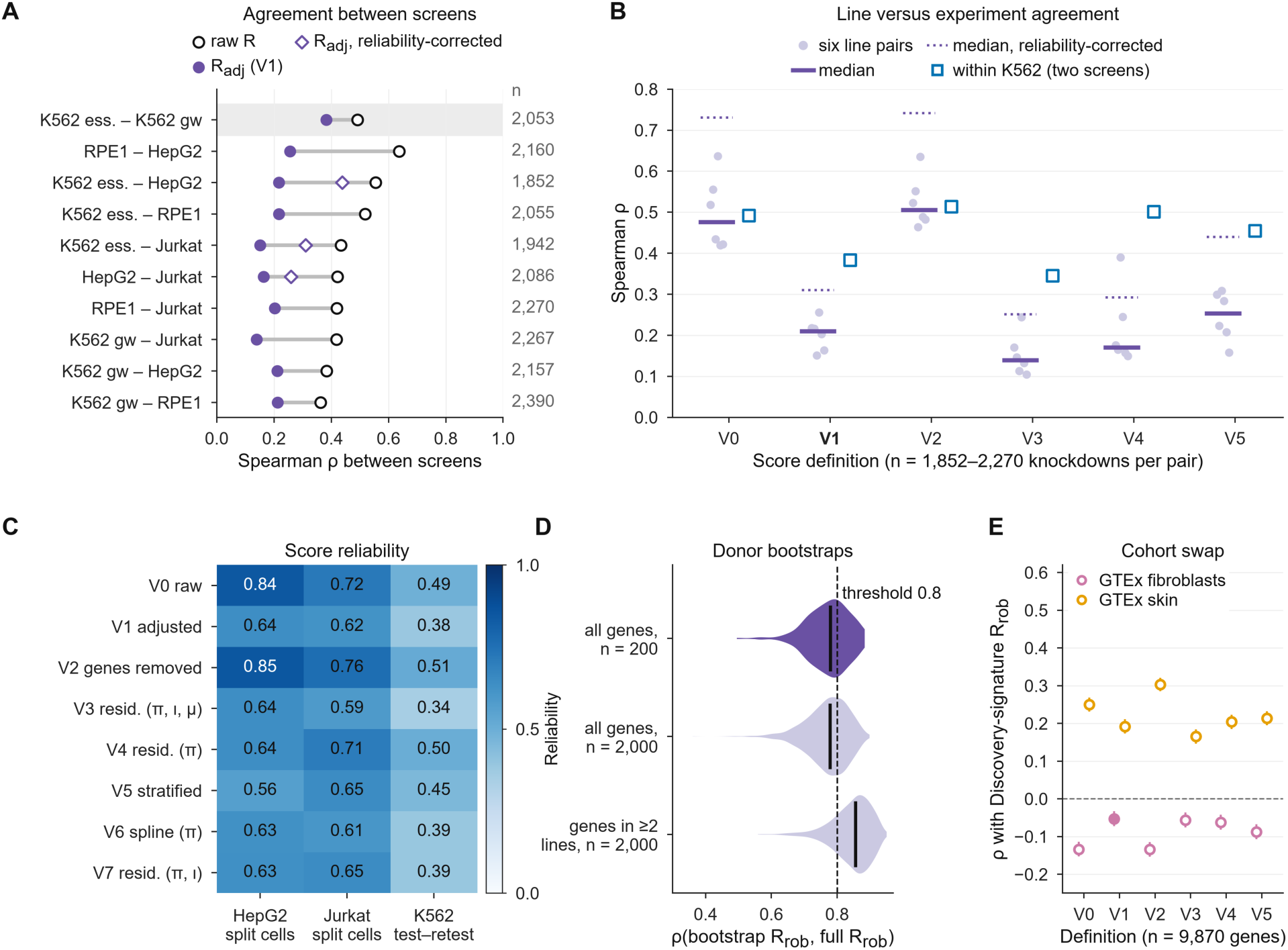
Reproducibility across lines, experiments, donors and cohorts. (A) Spearman ρ for all ten pairs of screens over shared knockdowns (n at right): raw R (open circles), R_adj_ (filled) and R_adj_ corrected for score reliability (diamonds). Shaded row, the within-line reference. (B) Between-line ρ for each score definition (light purple dots, the six pairs of lines; bar, median; dotted line, median corrected for reliability) and within-K562 ρ (squares) (*n* = 1,852–2,270 knockdowns per pair). (C) Split-cell (HepG2 and Jurkat) and test–retest (two K562 screens) reliability of each definition. (D) Correlation of R_rob_ from donor bootstraps with the full-cohort R_rob_, for all 9,870 genes and the 2,393 multi-line genes; dark violin, the test in Table 1 (H5a); n, bootstraps; solid lines, medians; dashed line, threshold 0.8. (E) Correlation of discovery-signature R_rob_ with R_rob_ from GTEx-derived signatures for V0 to V5 (*n* = 9,870 genes); error bars, 95% CIs; filled, the V1 GTEx-fibroblast test (H5b). Raw R in (A) tests H1a (Table 1); other panels are exploratory except as marked. See also Table S12.

Score reliability was limited (split-cell, Spearman–Brown corrected^36,37^: R_adj_ 0.64 in HepG2 and 0.62 in Jurkat; K562 test–retest 0.38; Figure 3C; Table S12). Corrected for reliability, adjusted agreement between K562 ess., HepG2 and Jurkat rose to 0.26–0.44 (upper estimates for pairs with K562) but stayed below raw agreement (0.54–0.86), so the adjusted signal is partly limited by noise and partly specific to each line (exploratory).

### Rankings change when the aging cohort changes

For the 7,477 of 9,870 genes measured only in the K562 genome-wide screen, R_rob_ equals R_adj_, so we report R_rob_ on all genes and on the 2,393 genes measured in at least two lines (Table S13). Across 200 donor bootstraps, the median rank correlation with the full-cohort ranking was 0.779, just below our threshold of 0.8 (Table 1). In 2,000 bootstraps (exploratory), a median 47% of top-5% candidates were kept (Figure 3D). On the multi-line genes the median was 0.856, so the stability failure is concentrated in genes measured in one screen.

Rankings did not transfer between cohorts. R_rob_ recomputed with the GTEx-derived signature correlated with the discovery ranking at ρ = −0.05 (one-sided *P* > 0.99), and only 4% of top-5% candidates were kept, against 47% between donor bootstraps. Because genes are not independent, the top-5% overlap is more useful than this P value. The result held on multi-line genes (ρ = −0.07; 3% kept) and for every definition (Figure 3E). A ranking from discovery donors in the GTEx age range (20 to 79) did not transfer either (5% kept, the chance rate; Table S3), but it was itself unstable, so this check cannot separate a cohort difference from noise. An exploratory GTEx skin-derived ranking agreed only weakly with the discovery ranking (ρ = 0.19; 11% of top-5% candidates shared, against 5% by chance). As genes are not independent and most come from one screen, we attach no P value and do not call this a replication.

### A protein network adds ranking signal but little accuracy

We asked whether a STRING network^38^ helps predict a held-out knockdown’s adjusted score (five-fold cross-validation within each screen; STAR Methods). The network predictor is the STRING-weighted mean score of high-confidence partners.

Comparators were degree-preserving randomized networks^39^, a constant, line transfer (the knockdown’s mean score in the other lines) and calibrated line transfer (a ridge model that rescales line transfer to the target screen).

The network beat ten degree-preserving randomizations on absolute error in 3 of 5 screens (RPE1, HepG2 and Jurkat; Table S14), so it passed this test (Table 1), but in the K562 genome-wide screen its error was 6.3% higher than the null (Figure 4A). Line transfer + network also beat uncalibrated line transfer in all five screens (Holm *P* < 0.001; Table 1), mostly through recalibration (see below).

**Figure 4.**
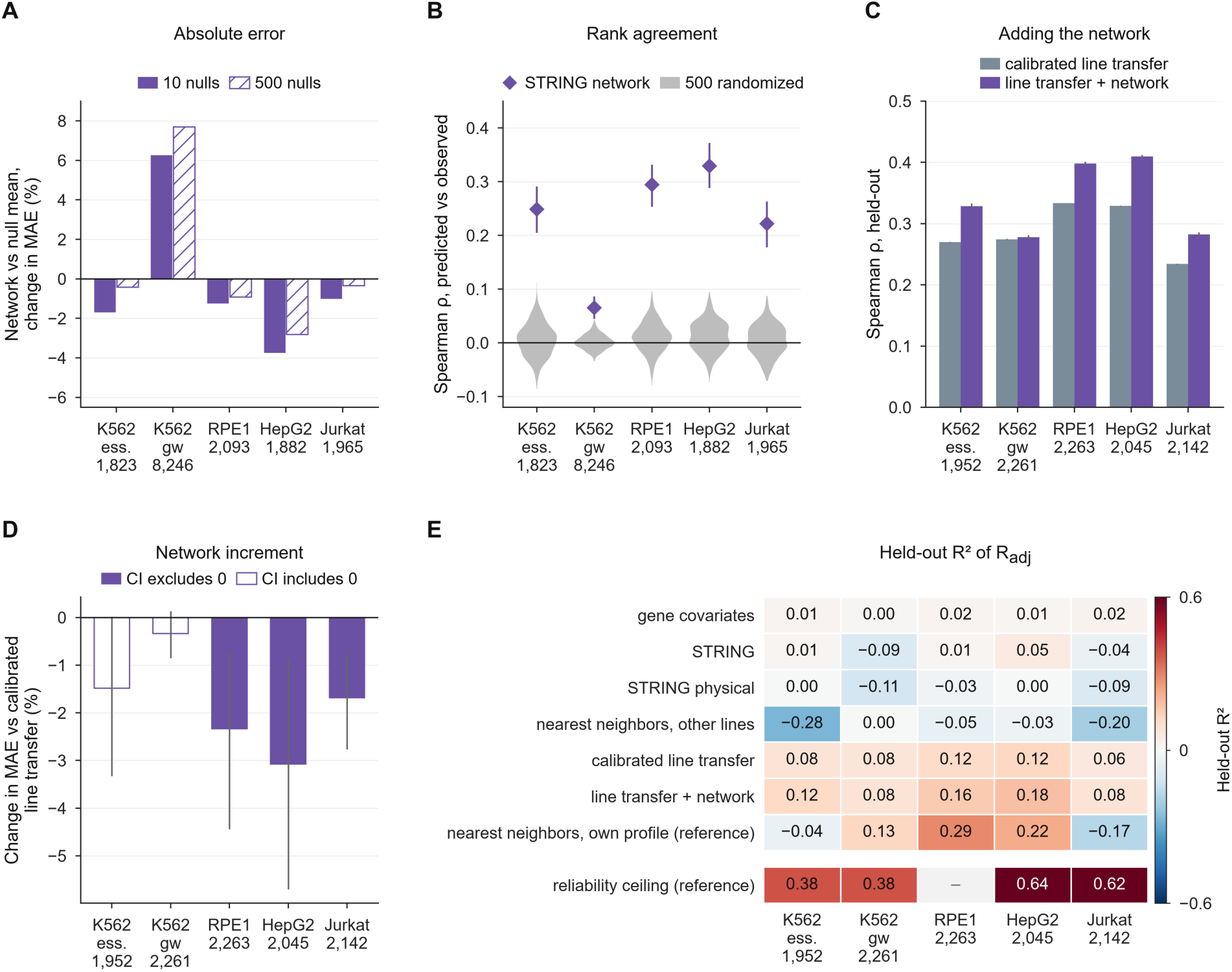
Incremental value of a protein interaction network. (A) Change in mean absolute error (MAE) of the network predictor relative to the mean of degree-preserving randomized networks: 10 nulls (filled) and 500 nulls (hatched); n, held-out knockdowns with a STRING node. (B) Spearman ρ between predicted and observed held-out R_adj_ for the network (diamonds; error bars, 95% bootstrap CIs) and 500 randomized networks (violins). (C) Spearman ρ of calibrated line transfer and line transfer + network (bars, mean over 10 cross-validation seeds; error bars, SD; n, knockdowns). (D) Change in MAE (%) of line transfer + network versus calibrated line transfer; error bars, 95% block-bootstrap CIs; open bars, CI includes zero. (E) Held-out R² of each predictor (rows) per screen (columns; mean over 10 seeds), with the reliability ceiling in the bottom row (–, not available for RPE1). The own-profile nearest-neighbor predictor is a reference only; R² = 0 for the constant predictor by design. The 10-null comparison in (A) tests H3a (Table 1); other panels are exploratory. See also Figure S8 and Table S14.

Against 500 randomizations (exploratory), the network’s ρ beat every null in all five screens (one-sided empirical *P* = 0.002; Figure 4B), but error fell significantly against their mean in only 2 of 5 screens. So passing depends on the number of nulls, whereas the rank advantage holds. Against calibrated line transfer, with every step refit inside training folds, line transfer + network raised ρ from 0.23–0.33 to 0.28–0.41 (Figure 4C) and changed mean absolute error (MAE) by −0.3% to −3.1% (Figure 4D); the block-bootstrap interval excluded zero in 3 of 5 screens. The network gives a small gain in some screens, not an improvement in every screen.

On held-out R², line transfer + network (0.08–0.18) was the best predictor that does not use the knockdown’s own profile; STRING alone and its physical subnetwork added little (Figures 4E and S8; Table S14). STRING ranked held-out knockdowns about as well as a nearest-neighbor predictor based on the gene’s response in other lines (ρ = 0.11–0.32 and 0.13–0.35). A nearest-neighbor reference that uses the knockdown’s own response on non-signature genes reached ρ = 0.36–0.52 (R² up to 0.29), so part of a knockdown’s score is predictable from the rest of its own profile.

### Agreement with held-out labels depends on the signature and matches a proliferation baseline

We tested scores against evidence never used in fitting (STAR Methods). CellAge^40,41^ labels genes whose manipulation induces or inhibits cellular senescence, whichever way they were manipulated, so knockdown of an inducer is expected to delay senescence. Of 370 inducers and 475 inhibitors, 257 and 394 had an R_rob_ score, almost all from the K562 genome-wide screen (Figure S9; Table S15). GenAge^42^ model-organism genes mapped to human orthologs through the Alliance of Genome Resources^43^ gave 942 anti-longevity genes (reduced function extends lifespan) and 548 pro-longevity genes. Knockdown of a senescence inducer or an anti-longevity gene was expected to score higher.

R_rob_ did not separate the labels (Table 1; Figure 5A): the CellAge AUROC was 0.54 (95% confidence interval, CI, 0.49 to 0.58; Holm *P* = 0.10) and the GenAge AUROC 0.51 (95% CI 0.47 to 0.55; Holm *P* = 0.27), and no other proliferation-adjusted definition did better (Figure 5B). The smallest AUROC detectable with 80% power was 0.56 for CellAge and 0.55 for GenAge, so these tests cannot rule out a small effect. In exploratory analyses, raw scores separated CellAge labels (AUROC 0.57; q = 0.017; Table S5) and pointed opposite to expectation for GenAge (0.44; q = 0.014), as did proliferation shift alone (CellAge 0.58, q = 0.011; GenAge 0.42, q = 0.0047). For CellAge, the raw score did not differ from proliferation shift (paired DeLong^44^ *P* = 0.57; Figure 5C) and added nothing once proliferation was modeled (Table S16). Adjustment did not clearly lower the CellAge AUROC (*P* = 0.095) and moved the GenAge AUROC toward chance (0.51; paired q < 0.001).

**Figure 5.**
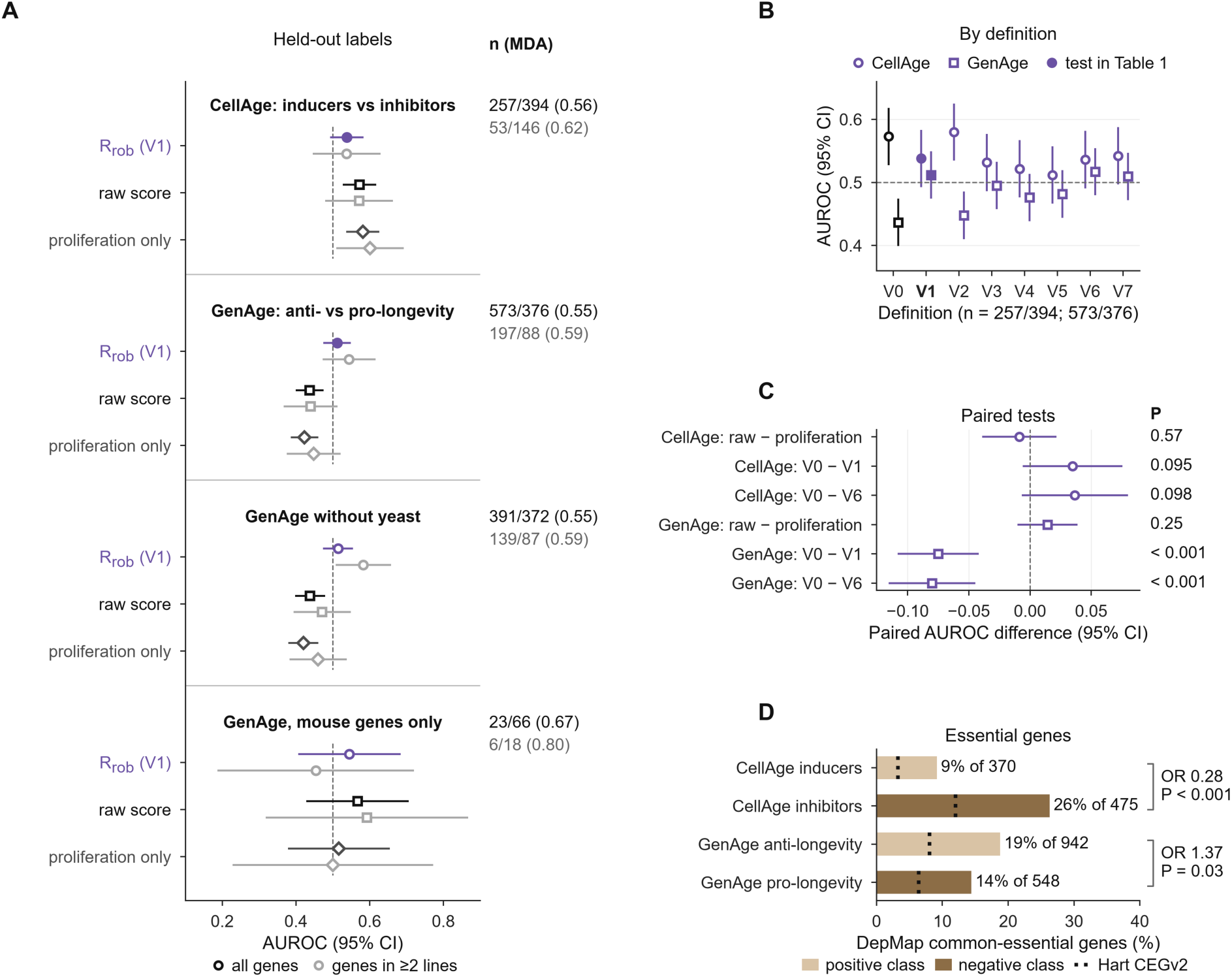
Held-out evidence. (A) AUROC for separating CellAge senescence inducers (positive class) from inhibitors and GenAge anti-longevity (positive class) from pro-longevity orthologs, by R_rob_ (V1; circles), the raw score (squares) and proliferation shift alone (diamonds); error bars, 95% Hanley–McNeil CIs^51^. Colors, score type (dark gray, proliferation only); light gray, multi-line genes. Dashed line, chance. n, positive/negative genes for all genes (black) and multi-line genes (gray), with the minimum detectable AUROC at 80% power (MDA) in parentheses. (B) AUROC (95% CI) of R_rob_ under each definition, all genes (*n* = 257/394 CellAge and 573/376 GenAge genes; V0 uses the raw score); circles, CellAge; squares, GenAge. (C) Paired AUROC differences with 95% CIs and two-sided DeLong P values (uncorrected; n as in B). (D) Fraction of each label class that is a DepMap 24Q2 common-essential gene (bars) or a Hart CEGv2 core-essential gene (dotted ticks); odds ratio (OR) and two-sided Fisher *P* (uncorrected; DepMap; n per class at the bar). In every panel, filled markers are the tests in Table 1 (V1 in A and B) and open markers are exploratory. See also Figure S9 and Tables S15, S16 and S17.

Label subsets did not show a proliferation-independent signal after correction (STAR Methods); the mouse-only GenAge set was too small to detect an AUROC below 0.67. With the GTEx-derived signature (exploratory), V1 separated CellAge labels at 0.58 (q = 0.0092; Table S5), equal to proliferation alone (paired DeLong *P* = 0.84), although V1 is adjusted for proliferation. On multi-line genes it also separated GenAge labels in the expected direction (0.625; q = 0.0092), where proliferation pointed the other way (0.45), but not on all genes (0.53; Figure S7; Table S11). This was the only score above chance for both label sets, so we treat it as a hypothesis for a new test. Label agreement depends on the signature and must be compared with a proliferation-only baseline every time.

CellAge labels are strongly enriched for genes that affect growth; GenAge labels only weakly. Of CellAge inhibitors, 26% are DepMap common essentials, against 9% of inducers (DepMap 24Q2^45,46^; Fisher *P* < 0.001; Figure 5D; Hart core-essential genes^47^: 12% versus 3%); for GenAge the split is 19% versus 14% (q = 0.086, not significant). As the two label sets point in opposite directions on proliferation, no score driven mainly by proliferation can agree with both. Senescence programs also differ by inducer^48^, and we know of no public dataset that measures senescence or another functional aging phenotype for these knockdowns in these lines; such data are the main validation gap. *CDKN1A* and *CDKN2A* are not targeted in any screen, and *RB1* moved from rank 128 (raw) to 1,937 (V1) (Table S17).

### Multi-objective ranking favors knockdowns with weak effects

Of the 2,393 multi-line genes, 16 were Pareto-optimal on five objectives: R_rob_, sign consistency, identity preservation, absence of proliferation gain and absence of stress induction (Figure S10; Table S18). A knockdown that changes little scores well on every safety objective, so 6 of the 16 had weak or unmeasured on-target knockdown. The top gene, *ANLN*, is a CellAge senescence inhibitor whose knockdown slows growth, so it scores well for a reason unrelated to rejuvenation. After quality filtering, 15 genes were on the front (STAR Methods). Every candidate is a hypothesis conditional on the discovery signature, not a rejuvenation target.

## DISCUSSION

Five of the nine main tests did not support our hypotheses (Table 1). Many findings came from exploratory analyses that need a new test.

We cannot say which cohort best reflects fibroblast aging in vivo. The reversal of the strongest age genes does not survive multiple-testing correction. Modeling GTEx batch removed most GTEx age genes, so we cannot tell technical structure from a real cohort difference.

Proliferation is a known confounder of cancer and aging signatures^26,27^, and here we measure it in perturbation screens. Because it is part of aging biology, we do not claim the adjusted score is correct.

After adjustment, the cross-line signal is partly line-specific and partly limited by noise, so rankings from one line may not transfer to another. Network information helped a little, consistent with network-propagation approaches^49^.

The held-out labels are also limited. CellAge labels are enriched for growth genes, and the two sets point in opposite directions on proliferation. A fair test needs functional aging read-outs for the same knockdowns in primary cells.

Our results do not contradict earlier screens^7,10,11^, which used other signatures, clocks, perturbation types and validation strategies; we did not rerun their methods.

## LIMITATIONS OF THE STUDY

First, and most important, none of the four lines is a fibroblast. All four have bypassed replicative senescence, and two (K562 and Jurkat) lack functional p53^50^. We therefore measure how knockdowns shift expression along a fibroblast aging axis in cancer and immortalized cells, not whether they rejuvenate an aged cell. Second, the libraries are dominated by essential genes; only the K562 genome-wide screen covers non-essential genes, and the held-out label tests rest mostly on that screen. Third, GTEx ages are binned, and culture passage is unavailable for both fibroblast cohorts, so cohort effects cannot be fully separated from aging. Fourth, we used two signatures; clocks from other tissues or from DNA methylation may behave differently. Fifth, this work used public data only and did not test any prediction experimentally. Sixth, no ranking passed every Box 1 check, including two halves of one GTEx cohort. The effect-size check compares independent cohorts with bootstraps that share donors, so it is strict by design; the stability threshold of 0.8 was fixed in advance. All sensitivity analyses came after the main results, and the analysis plan was written by authors who knew these public datasets well. Further, we pseudobulked HepG2 and Jurkat with the published normalization, which may differ in detail from the release used for the other screens. CRISPRi knockdown is not equivalent to pharmacological inhibition or to the loss-of-function alleles behind GenAge annotations. Sex is used only as a covariate and as a positive control.

## RESOURCE AVAILABILITY

### Lead contact

Requests for further information and resources should be directed to and will be fulfilled by the lead contact, Mohammad Ali Moni.

### Materials availability

This study did not generate new materials or new experimental data.

### Data and code availability

- This paper analyzes existing, publicly available data. Accession numbers, versions, access dates and checksums are listed in the key resources table and in the repository.
- All original code has been deposited at GitHub and Zenodo and is publicly available as of the date of publication (DOI in the key resources table). Gene-level scores, the signatures and the mapped labels are in Data S1 (Table S13). The HepG2 and Jurkat pseudobulk matrices will be deposited at Zenodo before publication (DOI in the key resources table).
- Any additional information required to reanalyze the data reported in this paper is available from the lead contact upon request.

## ACKNOWLEDGMENTS

We thank the authors of the Fleischer et al., Replogle et al. and Nadig et al. studies, the GTEx Consortium, and the STRING, MSigDB, Human Ageing Genomic Resources, DepMap, OncoKB and Alliance of Genome Resources teams. Icons in Figures 1A and 2A and in the graphical abstract: Servier Medical Art (smart.servier.com), CC BY 4.0 (fibroblasts, culture dish, skin, older man and woman, K562, HepG2 and Jurkat cells, mouse); dCas9 from CRISPR-Cas9 by Marcel Tisch (Bioicons, CC0; cut sites removed, KRAB domain added); network by Chenxin Li (Bioicons, CC BY 4.0); RPE1 cells, the worm, the cell-cycle wheel and the score bar drawn by us. This work was supported by the US National Science Foundation (award 2319522 to L.Z.).

## AUTHOR CONTRIBUTIONS

Conceptualization, M.K.; methodology, M.K. and S.A.D.; software, M.K. and S.A.D.; formal analysis, M.K.; validation, S.A.D., B.K.P. and A.A.N.; writing – original draft, M.K.; writing – review and editing, all authors; supervision, L.Z. and M.A.M.

## DECLARATION OF GENERATIVE AI AND AI-ASSISTED TECHNOLOGIES

During the preparation of this work, the authors used Claude (Opus 5.5; Anthropic) to write analysis code, produce figures, check references and edit the text. After using this tool, the authors reviewed and edited the content as needed and take full responsibility for the content of the publication.

## SUPPLEMENTAL INFORMATION

Document S1. Figures S1–S10 and Tables S1, S2, S6, S7, S8, S10, S11, S12, S15, S17, S18, S19 and S20.

Data S1. Tables S3, S4, S5, S9, S13, S14 and S16 (Excel workbook, one or more sheets per table):

- Table S3. Additional statistics reported in the text (power checks, random signatures, hidden factors, top-k sweep, split halves, reliability and bootstrap summaries), related to Figures 1 to 3
- Table S4. Cross-cohort agreement for every cohort specification, related to Figure 1
- Table S5. All exploratory tests with a P value, with family sizes and corrected values, related to STAR Methods
- Table S9. Score definitions per screen, related to Figure 2
- Table S13. Gene-level scores, the age signatures and the mapped labels, related to STAR Methods
- Table S14. Network prediction: main tests, in-fold baselines and block bootstrap, related to Figure 4
- Table S16. Held-out evidence tests for every score definition and label subset, related to Figure 5

## BOX 1.

**Checklist for transcriptomic rejuvenation screens**

In every rule, corrected P means Holm-adjusted P for tests fixed before the study and Benjamini–Hochberg q for other tests, within each check.

1. Cohorts. Report rankings under at least two independent aging cohorts. Judge agreement among the strongest age genes against a permutation null, check with a shared effect planted into the real data that the test could detect agreement, and run a positive control (such as sex) through the same pipeline. Pass: top-200 ρ above the 97.5th percentile of the permuted-age null under the main model; a shared effect added to the real data, with planted |t| no larger than the median |t| of the observed top-k genes, recovered in at least 80% of runs; positive control above its own null (corrected *P* < 0.05).
2. Programs. Report the correlation of the raw score with proliferation, stress and response size in every screen, and report rankings with and without proliferation adjustment. Compare the real signature with random signatures from the same gene pool. Flag: |ρ| with proliferation ≥ 0.3 in any screen, or a real signature that tracks proliferation more than 95% of random signatures; the link is then specific to the signature and must be reported with the ranking.
3. Replicates. Compare agreement between two experiments in one line with agreement between lines, report the two side by side, and report score reliability for every definition. Pass: reliability of the adjusted score ≥ 0.5 in every screen, and between-line agreement of the adjusted score above the random-signature null.
4. Labels. Validate against evidence not used for model selection, next to a proliferation-only baseline. Pass: AUROC above 0.5 (corrected *P* < 0.05) and above the proliferation-only AUROC (paired DeLong, corrected *P* < 0.05), for each label set.
5. Power. Report the minimum detectable AUROC for every label test; a test with a minimum detectable AUROC above 0.60 is reported as uninformative.
6. Effect sizes. Judge agreement between rankings by top-k overlap and effect size, not by P values over thousands of genes. Pass: top-5% overlap between cohorts at least half the overlap between donor bootstraps of one cohort.
7. Knockdown quality. Report the share of top-ranked or Pareto-front genes with weak or unmeasured on-target knockdown before filtering, and rank only after filtering on on-target knockdown and response size.
8. Stability. Pass: median donor-bootstrap ρ of the ranking ≥ 0.8.

Applied to the discovery signature, the ranking fails the cohort, replicate, label, effect-size and stability checks (stability on all genes only) and is flagged for proliferation (Table S20). Under these rules, its top candidates are hypotheses only and should not be used to choose rejuvenation targets. The GTEx-derived signature fails the same checks (Table S20). As a positive example, we split GTEx fibroblasts into two random halves of 326 donors (Table S3). The halves agreed on the strongest age genes (top-200 ρ = 0.71 against a null 97.5th percentile of 0.58) and agreed more on rankings than the two cohorts did in every one of 20 splits (ρ = 0.11 to 0.52 against −0.05). Rankings within one cohort are unstable (median donor-bootstrap ρ = 0.43 for one half, so the halves also fail the stability and effect-size checks), but swapping cohorts is worse. Each half has more donors than the discovery cohort (326 against 111), which favors the halves. The stability rule and the AUROC-above-0.5 part of the label rule follow tests fixed before this study (Table 1); the comparison with the proliferation-only baseline and the other thresholds are our suggestions, set after this study, and untested on other screens.

## STAR★METHODS

### KEY RESOURCES TABLE

| REAGENT or RESOURCE | SOURCE | IDENTIFIER |
|---|---|---|
| **Deposited data** |  |  |
| Primary dermal fibroblast RNA-seq (FPKM, fragments per kilobase per million reads) and series matrices | Fleischer et al.<sup>21</sup> | GEO: GSE113957 |
| GTEx v10 gene TPM (transcripts per million; cultured fibroblasts; skin, not sun-exposed) and open-access annotations | GTEx Consortium<sup>22,30</sup> | https://gtexportal.org; release v10 |
| K562 essential-gene, K562 genome-wide and RPE1 CRISPRi Perturb-seq pseudobulk | Replogle et al.<sup>17</sup> | Figshare: 20029387 |
| HepG2 and Jurkat CRISPRi Perturb-seq raw counts | Nadig et al.<sup>18</sup> | GEO: GSE264667 |
| STRING v12.0 human protein links and physical subnetwork | Szklarczyk et al.<sup>38</sup> | https://string-db.org; v12.0; RRID: SCR_005223 |
| MSigDB Hallmark gene sets (via Enrichr library MSigDB_Hallmark_2020) | Liberzon et al.<sup>32</sup>; Kuleshov et al.<sup>52</sup> | https://maayanlab.cloud/Enrichr; RRID: SCR_016863 |
| CellAge (cellAge.zip, 866 genes; web table with manipulation methods) and GenAge model organisms | Avelar et al.<sup>40</sup>; de Magalhães et al.<sup>42</sup> | https://genomics.senescence.info |
| Core-essential genes (CEGv2) | Hart et al.<sup>47</sup> | https://doi.org/10.1534/g3.117.041277 |
| CRISPR-inferred common-essential genes, DepMap 24Q2 | Broad DepMap<sup>45,46</sup> | https://doi.org/10.25452/figshare.plus.25880521.v1 |
| OncoKB cancer gene list | Chakravarty et al.<sup>53</sup> | https://www.oncokb.org; accessed 12 October 2026 |
| Combined orthology (ORTHOLOGY-ALLIANCE_COMBINED) | Alliance of Genome Resources<sup>43</sup> | https://www.alliancegenome.org; accessed 12 October 2026 |
| TRACE code, results and figure source data | This paper | Zenodo: https://doi.org/10.5281/zenodo.23763631 |
| HepG2 and Jurkat pseudobulk expression shifts | This paper | GitHub: https://github.com/SaiKawserKazi/TRACE; Zenodo: https://doi.org/10.5281/zenodo.23763631 |
| **Experimental models: Cell lines** |  |  |
| K562 (female) | Replogle et al.<sup>17</sup> | RRID: CVCL_0004 |
| hTERT (telomerase reverse transcriptase)-immortalized RPE-1 (female) | Replogle et al.<sup>17</sup> | RRID: CVCL_4388 |
| HepG2 (male) | Nadig et al.<sup>18</sup> | RRID: CVCL_0027 |
| Jurkat (male) | Nadig et al.<sup>18</sup> | RRID: CVCL_0065 |
| **Software and algorithms** |  |  |
| Python 3.13.7 | Python Software Foundation | https://www.python.org; RRID: SCR_008394 |
| NumPy 2.5.2; SciPy 1.18.1 | Harris et al.<sup>54</sup>; Virtanen et al.<sup>55</sup> | RRID: SCR_008633; RRID: SCR_008058 |
| pandas 3.0.5 | The pandas development team | https://pandas.pydata.org; RRID: SCR_018214 |
| statsmodels 0.15.0; patsy 1.0.3 | The statsmodels developers | https://www.statsmodels.org; RRID: SCR_016074 |
| scikit-learn 1.9.1 | Pedregosa et al.<sup>56</sup> | RRID: SCR_002577 |
| NetworkX 3.6.1 | The NetworkX developers | https://networkx.org; RRID: SCR_016864 |
| anndata 0.13.4; h5py 3.16.0; openpyxl 3.1.5 | The scverse, h5py and openpyxl developers | https://anndata.readthedocs.io; https://www.h5py.org; https://openpyxl.readthedocs.io |
| Matplotlib 3.11.1 | Hunter<sup>57</sup> | RRID: SCR_008624 |
| TRACE pipeline and figures | This paper | Zenodo: https://doi.org/10.5281/zenodo.23763631 |

### EXPERIMENTAL MODEL AND STUDY PARTICIPANT DETAILS

This study generated no new experimental data and involved no new human participants. It analyzed published, de-identified data: dermal fibroblasts from healthy donors^21^, GTEx post-mortem and organ donors (open-access tier)^22^, and CRISPRi Perturb-seq of K562 (chronic myeloid leukemia; female), RPE1 (telomerase-immortalized retinal pigment epithelium^58^; female), HepG2 (hepatoblastoma; male) and Jurkat (T cell acute lymphoblastic leukemia; male) cells^17,18^. Cell line sex and *TP53* status are as listed in Cellosaurus^50^ (RRIDs in the key resources table). The cell lines were used as published data only; we did not culture or authenticate them. All human data are public and de-identified and were collected under the approvals of the original studies, so no new approval was needed. Donor age and sex per cohort are summarized in Table S2 (discovery 111 donors, 29 female, aged 19–96 years; GTEx fibroblasts 652 donors, 216 female; GTEx skin 651 donors, 206 female; ancestry is available for the discovery cohort only). Sex enters the analyses as a covariate and as a positive control; no sex-stratified analysis was done.

### METHOD DETAILS

#### Study design

We fixed five hypotheses and their tests before scoring any knockdown (protocol and timestamp in the repository). All later analyses are labeled exploratory. The protocol file was committed on 12 October 2026 (commit 1eb8681, tag protocol-v1.0) and is unchanged since; its SHA-256 is 01a573574e8225e5ba68ddd2757b81dcd765083274d9b3290b8a1bb0ca879fa7. The protocol, the first run and the first deviations were made on the same day; the commit history fixes their order. The protocol was not deposited with a third-party registry, so its freeze time rests on version control, and we call the analyses pre-specified rather than pre-registered. Deviations from the protocol and the dated list of exploratory analyses are in Table S19. In brief, orthologs came from the Alliance of Genome Resources because the Ensembl BioMart service was unavailable; the two K562 screens count as one line for R_rob_, sign consistency and the Pareto ranking, and the K562 essential-gene screen stands for K562 in H1; HepG2 and Jurkat were pseudobulked by us; the H3 ridge penalty was set to λ = 1; a file-size check silently skipped the yeast ortholog table in the first H4b run, after the first H4 results were seen, and the corrected run changed the H4b AUROC from 0.52 to 0.51; and an error in an exploratory skin model (ischemic time in G0) was corrected without affecting any pre-specified result. No pre-specified outcome changed, but the 500-null analysis showed that the H3a result is fragile.

#### Aging cohorts and signatures

FPKM values (GSE113957) were summed per gene symbol. Donor age, sex and ethnicity were parsed from sample titles, and platform, biopsy site and repository cell ID were taken from the two GEO platform series matrices. From the 133 healthy donors in GSE113957 (143 samples, including 10 progeria samples), we excluded the 22 donors younger than 18, leaving 111 donors aged 19–96 years by the GEO age field. AG denotes the NIA Aging Cell Repository and GM the National Institute of General Medical Sciences (NIGMS) Human Genetic Cell Repository. GTEx v10 gene TPMs were summed per symbol and combined with open-access subject and sample annotations. Age-bin midpoints were used, and one sample was kept per donor. Expression was log2(x + 1) transformed. For genes with mean log2 expression ≥ 1, expression ∼ age + covariates was fitted by ordinary least squares, with categorical covariates dummy-coded and numeric covariates median-imputed. P values were corrected with the Benjamini–Hochberg procedure. The pre-specified model used sex only. The signature comprised up to 250 up- and 250 down-regulated genes with FDR < 0.05, ranked by t. If fewer than 50 genes passed in either direction, the code used the top 250 up and down genes by t without the FDR filter; among the point estimates this applied only to F8, which had no FDR < 0.05 genes, and the number of bootstrap replicates that used it was not recorded. Hallmark enrichment used one-sided hypergeometric tests against expressed genes. The GEO sample titles give ages of 19 to 96 years for the retained donors, whereas the original study reports 1 to 94 years. We used the GEO values; limiting the analysis to donors aged 20 to 79 removes this uncertainty. The variance inflation factor of age is 1.43 in a discovery model with sex, platform, repository and biopsy site. In GTEx, age overlaps less with nucleic-acid isolation batch in fibroblasts (variance inflation factor 1.18 with batch alone and 1.41 in the full GTEx batch model, G2) and more in skin (1.64 and 2.05; Table S3). So age and repository overlap, but collinearity is modest. Thirteen discovery samples, all from donors younger than 60, were sequenced on a different platform.

Cohort sensitivity (exploratory). There were ten discovery specifications: F0 pre-specified; F1 ages 20–79; F2 + platform, repository and biopsy site; F3 = F1 + F2; F4 NextSeq only; F5 arm biopsies only; F6 and F7 expression filters 0.5 and 2; F8 without AG donors aged 80 or older; and F9 + five hidden factors. There were four replication specifications: G0 age + sex; G1 + center, batch type, RIN and Hardy scale (skin also ischemic time); G2 = G1 + nucleic-acid isolation batch; and G3 age + sex + five hidden factors. Hidden factors were the first five principal components of the residual expression after the age + sex model; we used five factors without tuning and report what they track. Because these residuals are orthogonal to age, the factors leave each gene’s age coefficient unchanged and change only its standard error. As a check (F9b), we also used the first five principal components after a sex-only model, which do not protect age, with their own permuted-age null. CIs came from 2,000 donor-level bootstraps of each cohort, refitting the model and re-deriving the signature each time; the same donor resample was used for all models of a cohort within a replicate. Intervals are bias-corrected percentile intervals (no acceleration term). For the 200 strongest genes, bootstrap estimates sit near the ends of their CIs because each replicate re-derives the 200 strongest genes. Split-half references used 500 random halvings per cohort. The permuted-age null shuffled age within each cohort 1,000 times with sex and covariates fixed, for every discovery specification against G0. Empirical two-sided P values are (1 + number of nulls with |ρ| ≥ |observed|)/(1 + 1,000). The sex control repeated the same models with sex as the variable and reported Spearman ρ of t-statistics over all genes, autosomal genes, X and Y genes, and the 200 genes with the largest discovery effect, with a permuted-sex null (1,000 permutations). In the planted-effect check, age was permuted in both cohorts and a shared effect was added to 500 random expressed genes: k × SD(gene) × z(age) with a random sign shared by both cohorts, for k = 0.2 and 0.41 (200 replicates each). The planted effect touched 500 genes; power for a shared effect spread over more genes was not tested. Effects were planted after age was permuted, so they did not compete with real age effects. We repeated the check with the effect added to the real data of both cohorts (k = 0.38 and 0.57, giving a planted part alone of median |t| about 4 and 6; 200 replicates each). The top-k sweep, the sign agreement of signature genes and F9b were tested against the same 1,000 permuted age labelings as F0, with the signature rebuilt from each permuted discovery fit; empirical P values for sign agreement are taken about the null median. The agreement of GTEx fibroblasts with GTEx skin was tested against 1,000 permutations of age within each tissue. For G2, a further null permuted age within nucleic-acid isolation batch in GTEx (donors in single-donor batches, 28 of 72 batches in fibroblasts and 73 of 243 in skin, keep their age) and across donors in the discovery cohort, 1,000 times, refitting G2 each time. Batches held on average 9.1 donors in fibroblasts and 2.7 in skin, so in skin this null keeps most of the age structure and is weak. For the Box 1 cohort check of the GTEx-derived signature, the 200 strongest GTEx-fibroblast age genes were compared with the discovery cohort under the same 1,000 permuted age labelings as F0. For the like-for-like check, the ranking was rebuilt from discovery donors aged 20 to 79 (signature rule as above) and compared with the GTEx-fibroblast ranking without a P value; 100 donor bootstraps of this subset gave the reference. For the Box 1 positive example, GTEx fibroblasts were split into two random halves; for one split we applied the cohort check (200 permuted age labelings in both halves), the effect-size check and the stability check (100 donor bootstraps of one half), and 20 splits gave the spread of the point estimates. The power-matched check drew 1,000 random 74-donor subsets of the 111 donors. Age prediction used an elastic net (scikit-learn^56^, l1 ratio 0.5, penalty chosen by inner five-fold cross-validation) on genes expressed in both the training and the test cohort; the number of genes with a non-zero coefficient is given in Table S8.

Additional cohort results (exploratory). Under 1,000 permuted age labelings of the discovery cohort, the 95th percentile of the number of FDR < 0.05 age genes was 0 and the 99th percentile 4, but one permutation gave 3,178, because correlated genes can produce rare large false hits. The 1,000 random 74-donor subsets gave a median of 5,128 age genes (95% range 2,618 to 6,596). The agreement of GTEx fibroblasts with GTEx skin had a permuted-age 95% range of −0.22 to 0.22. Two random halves of one cohort agreed at a median ρ of 0.84 (discovery), 0.76 (GTEx fibroblasts) and 0.84 (GTEx skin) over 500 random splits. Removing cell-cycle genes changed the genome-wide correlations little (ρ = −0.25 with GTEx fibroblasts and 0.16 with skin). With GTEx skin, top-k ρ was positive but fell from 0.26 to 0.08 as k grew. The six discovery specifications in which the top-200 reversal fell inside the permuted-age null were adjustment for five hidden factors (−0.08) or for platform, repository and biopsy site (−0.19), ages 20 to 79 only, without (−0.33) or with these covariates (−0.14), arm biopsies only (−0.41), and no AG donors aged 80 or older (−0.37; Table S4); these subsets are smaller and have wider nulls. The reversal stayed outside the null with NextSeq samples only (−0.61) and with either expression filter (−0.56 and −0.55). In GTEx fibroblasts, 22.1% of signature genes moved in the same direction, against a permuted-age 95% range of 23.3% to 76.8% (permutation *P* = 0.050, uncorrected; Figure S2); the range is wide because correlated genes move together. The median age t-statistic of E2F-target and G2M-checkpoint genes was −4.3 and −4.2 in the discovery cohort, but 2.0 and 1.8 in GTEx fibroblasts. In the sex control, agreement among the 200 most sex-biased autosomal genes was 0.34 with GTEx fibroblasts (permuted-sex *P* = 0.14) and −0.18 with skin (*P* = 0.25), and over all autosomal genes 0.17 and −0.13 (Table S7). The real 200 strongest discovery age genes all had |t| ≥ 5.9 (median 6.2). When the shared effect was added to the real data, planted genes made up a median of 67% and 96% of the 200 strongest discovery genes, and the median top-200 ρ with GTEx fibroblasts was 0.78 and 0.73 (Table S3). The planted effect was linear in age. Across the 80 specification combinations, genome-wide ρ ranged from −0.31 to 0.26. Under G2, top-200 values ranged from −0.18 to 0.13. The null that permuted age within nucleic-acid isolation batch gave a 95% range of −0.44 to 0.44 for the G2 value of F0 (−0.17), although G2 kept more age genes than 99.5% of permuted fits (515 against a median of 0). Hidden factors: factor 1 was uncorrelated with age (ρ = −0.04) and enriched for Myc targets V2 genes (q = 0.0048). Factors 2 and 3 tracked repository (area under the receiver operating characteristic curve, AUROC, for AG 0.30 and 0.32), and factor 3 tracked platform (AUROC for NextSeq 0.70). Because these factors do not change age effect sizes, adjusting for them (F9) changed only the standard errors. This reordered the genes by t, so a different set of genes reached the top 200 (29 of 200 shared), and among them the reversal was close to zero (−0.08); on the original 200 genes the F9 t-statistics gave −0.61. Of the sex-only factors (F9b), the first correlated with age (ρ = 0.38), and the second and third tracked repository (AUROC for AG 0.30 for both). Adjusting for them still left the reversal outside its permuted-age null (−0.40; permutation *P* = 0.019, uncorrected), so the reversal is not explained by the main hidden expression programs of the discovery cohort; with these factors, agreement with skin also turned negative (−0.31; *P* = 0.028; Table S3). Age prediction: within-cohort cross-validated correlation was 0.54–0.72, whereas across cohorts it was −0.09 to 0.29 (Table S8); within-cohort prediction used five-fold cross-validation, and its CIs used the Fisher z transformation. In the Box 1 positive example, the two GTEx halves kept 7% to 31% of top-5% candidates over the 20 splits, against 4% between the two cohorts.

#### Perturbation data

Pseudobulk shifts for the K562 essential-gene, K562 genome-wide and RPE1 screens were taken from Replogle et al.^17^: per-gene expression z-scored against non-targeting cells of the same gem group and averaged per perturbation. For HepG2 and Jurkat^18^, raw counts were scaled to the median non-targeting library size, z-scored per gene against non-targeting cells of the same gem group, and averaged per target gene, as planned for targets with at least 20 cells, restricted to genes measured in the Replogle release. Median on-target knockdown was comparable (HepG2 z = −0.73; K562 −0.71). Guides targeting one gene were averaged, and genes with non-finite values were removed. Non-targeting entries served as negative controls; for HepG2 and Jurkat these were 100 random groups of non-targeting cells, and size-matched groups (with the cell count of a typical knockdown) were added for the reliability analysis.

#### Score definitions

For perturbation p in screen c, with signature S and age t-statistics a, the raw score excludes the target gene p:

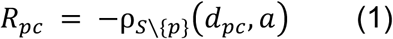

where ρ is the Spearman correlation over the genes in S other than p. Program shifts are mean z-shifts over gene sets: proliferation π (Hallmark E2F targets ∪ G2M checkpoint), stress σ (TNF-α signaling via NF-κB ∪ inflammatory response ∪ p53 pathway) and identity ι; response size μ is the mean absolute z-shift. The target gene is excluded from R but not from π, as in the protocol; one gene among about 300 has a negligible effect. Identity genes are the 100 genes most specific to a line: log-normalized gene means of each line minus the mean of the other lines, among genes above that line’s median expression (Table S19). For HepG2 and Jurkat the means come from non-targeting cells only; for K562 and RPE1 they are the gene means distributed with the Replogle release, which we could not confirm to be computed on control cells only. V1 is the residual of the score computed on S′ (S minus Hallmark E2F-target, G2M-checkpoint, mitotic-spindle and MYC-target genes) after least-squares regression on (1, π, ι, μ) across the knockdowns of a screen, in the spirit of growth-rate corrections^34^. V2 uses S′ without residualization, V3 residualizes the raw score on (π, ι, μ), V4 on π only, and V5 is (the percentile rank of R within its π quintile) − 0.5. V6 is a B-spline residual on π (five degrees of freedom), and V7 a residual on (π, ι) without μ. A plain up-minus-down score correlated with R at ρ = 0.93–0.98 (Table S9). The robust score is

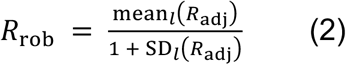

over lines l, with the two K562 screens averaged into one line and the population SD, so that SD = 0 and R_rob_ = R_adj_ for genes measured in one line. Sign consistency is computed over lines in the same way. In the reliability analysis, the cells of each knockdown were split into two random halves 20 times and the scores of the halves were correlated (Spearman–Brown corrected); for K562, the two screens served as test and retest. Disattenuated ρ is ρ divided by the square root of the product of the two reliabilities. Distance correlation and mutual information with π are given per screen and definition in Table S9. For pairs with K562, reliability is test–retest between two screens, which is lower than split-cell reliability, so the corrected values for these pairs are upper estimates; test–retest reliability is the same quantity as the within-K562 agreement. CIs for the cohort comparison of rankings (Figure 3E) used the Fisher z transformation.

Additional score, replicate and ranking results (exploratory). The screens held 2,057 (K562 ess.), 9,866 (K562 gw), 2,393 (RPE1), 2,160 (HepG2) and 2,270 (Jurkat) knockdowns, each measuring 363–399 signature genes. Non-targeting entries had narrower score distributions than knockdowns (SD 0.10–0.15 versus 0.13–0.22; Figure S3). In RPE1 and HepG2 the whole knockdown distribution shifted to about −0.5, consistent with most essential-gene knockdowns lowering cell-cycle genes. When proliferation was measured in one random half of each knockdown’s cells and R in the other, ρ fell from 0.89 to 0.68 in HepG2 and from 0.62 to 0.37 in Jurkat. The remaining non-linear dependence on π, measured by distance correlation, was 0.04–0.10 for V6. Per screen, top-5% overlap between definitions was lower than when pooled (Table S9); V6 and V7 shared 79 and 98 of the 119 top-5% genes with V1. The raw top 5% had a mean proliferation shift of 0.018, against −0.055 for V1 and −0.069 for all knockdowns. The GTEx-derived signature was enriched for IL-6/JAK/STAT3, TNF-α signaling via NF-κB and inflammatory-response genes in its down arm; its raw scores tracked stress (0.04–0.31) and response size (0.00–0.31) weakly and proliferation inversely (−0.39 to 0.07). V0 and V2, the two definitions that keep the proliferation component, are the only ones with no gap between within-line and between-line agreement. Split-cell reliability of the raw score was 0.84 (HepG2) and 0.72 (Jurkat), and K562 test–retest reliability of the raw score was 0.49; RPE1 reliability was not computed because we used only the pseudobulk RPE1 release. The 2,393 genes measured in at least two lines are the genes of the RPE1 library. Between cohorts, R_rob_ agreement was ρ = −0.13 to −0.05 for V0 to V5. The ranking from discovery donors aged 20 to 79 used 74 donors and 198 age genes and had a median donor-bootstrap ρ of 0.47. The GTEx skin-derived ranking correlated with the discovery ranking at ρ = 0.19 over 9,870 genes (0.17–0.30 across definitions), which explains about 4% of rank variance; 11% of the discovery top-5% candidates stayed in the skin-derived top 5%.

#### Random signatures

Each of 100 random signatures kept the 434 non-proliferation genes’ real age t-values and assigned them to genes drawn from a pool that excludes all proliferation genes, matched in bins of mean expression and variance (99.8% of draws from the exact bin). They were run through H1, H2 and H4 and compared with the real signature after its proliferation genes were removed (V2 input), so that neither side contains cell-cycle genes. One-sided empirical P values are (1 + number of random signatures at or above the real value)/(1 + 100). For K562 ess., the real signature gave 0.45, against a null median of 0.10 and maximum of 0.42. For adjusted cross-line agreement, the real signature gave 0.21 against a random-signature maximum of 0.18. For the adjusted CellAge AUROC, the empirical *P* was 0.040, with a null median of 0.489. The GTEx-derived signature had 3.6% cell-cycle genes in its down arm, against 23.2% for the discovery signature, and its raw scores correlated negatively with the oxidative-phosphorylation shift (−0.30 to −0.03), as expected for a program in its up arm.

#### Network prediction

For each screen, knockdowns with a STRING node were split into five folds (seed 0). The network predictor is the STRING-score-weighted mean R_adj_ of training-fold partners (combined score ≥ 700). Nulls are ten double-edge-swap randomizations (2|E| swaps, seeds 0–9, edge weights permuted). Degree preservation was checked for these ten nulls and for all 500 exploratory nulls (seeds 1000–1499). Line transfer is the mean R_adj_ of the same knockdown in the other lines; the line-transfer feature never uses a screen from the same cell line, so for K562 essential it excludes K562 genome-wide, and the reverse. The calibrated transfer and combined models are ridge regressions (λ = 1) fitted in training folds on line transfer alone and on line transfer plus network. In the exploratory in-fold analysis, residualization, quintile bounds and the ridge penalty (inner cross-validation over λ from 10^−6^ to 10^3^) were fitted on training folds only, ten cross-validation seeds were averaged, and all predictors were evaluated on one common subset per screen. Added baselines were gene covariates, the STRING physical subnetwork, a nearest-neighbor predictor (k = 20) that finds neighbors by the gene’s response profile in the other lines, the same predictor using the knockdown’s own response on non-signature genes (a reference), and proliferation shift as a predictor of the raw score. The block bootstrap resampled 100 clusters from k-means clustering of knockdown response profiles. Metrics are MAE, Spearman ρ and R² over held-out knockdowns, aggregated per screen. The H3 target R_adj_ is residualized on all knockdowns, test folds included, as pre-specified; the in-fold analysis removes this. The high-confidence network had 16,201 genes and 236,930 edges. In the 500 exploratory randomizations, every node degree was preserved and 96% of edges were rewired; the null 97.5th percentile of ρ was ≤ 0.07 in every screen. In the K562 genome-wide screen the network’s ρ was 0.07 over the 8,246 knockdowns of the original folds, against a null 97.5th percentile of 0.03. In the pre-specified H3b test, MAE fell from 0.0758–0.0825 (uncalibrated line transfer) to 0.0665–0.0755 (line transfer + network). Against calibrated line transfer, adding the network changed MAE by −1.5%, −0.3%, −2.3%, −3.1% and −1.7% in K562 ess., K562 gw, RPE1, HepG2 and Jurkat, with almost no change in Spearman ρ in K562 gw; the block-bootstrap CIs used 2,000 resamples. The inner cross-validation chose the smallest ridge penalty in 60–100% of folds, so these models were effectively unpenalized. On the common subset, held-out R² was −0.09 to 0.05 for the STRING network alone and −0.11 to 0.00 for its physical-interaction subnetwork, and −0.17 to 0.29 for the own-profile nearest-neighbor reference; the K562 genome-wide STRING ρ (0.11) was computed over 2,261 knockdowns, a different gene set from the 8,246 above. With 500 nulls, an empirical P of 0.002 is the smallest attainable. Proliferation shift alone predicted the raw score at ρ = 0.47–0.90 in held-out folds (common subset; Table S14).

#### Evidence labels

CellAge labels were assigned per gene from the distributed senescence effect (inducer or inhibitor), which describes the gene’s own effect whichever way it was manipulated; genes with conflicting records were dropped. The CellAge web table lists the manipulation method of each record, which we used for the loss-of-function subset. GenAge model-organism genes with Pro- or Anti-Longevity labels were mapped to human orthologs through Alliance pairs supported by at least three algorithms and best-scoring in at least one direction. Human labels were assigned by majority over organisms, and ties were dropped. Label counts and coverage are in Table S15; 400 of the 911 yeast genes carry a longevity label (most others are annotated as necessary for fitness), and 246 of these map to human orthologs. The CellAge release downloaded on 12 October 2026 lists 866 genes, more than the 279-gene build described in the 2020 paper; of the 257 inducers and 394 inhibitors with a score, 257 and 393 came from the K562 genome-wide screen (one inhibitor was measured only outside it). For GenAge, 49 ties were dropped after majority voting; worms contributed the most anti-longevity genes (578 worm genes, against 349 yeast genes, both before ortholog mapping).

Additional label results (exploratory). The 95% CIs were 0.53 to 0.62 for the raw-score CellAge AUROC (0.57), 0.40 to 0.47 for the raw-score GenAge AUROC (0.44), 0.54 to 0.62 for proliferation shift with CellAge (0.58) and 0.39 to 0.46 with GenAge (0.42). Across V1 and V3 to V7, AUROCs were 0.51 to 0.54 for CellAge and 0.48 to 0.52 for GenAge. For CellAge, the paired difference between the raw score and proliferation shift was −0.009 (95% CI −0.039 to 0.022); in a logistic model with proliferation, the raw-score coefficient was 0.03 (95% CI −0.22 to 0.29; likelihood-ratio *P* = 0.79); and the AUROC fell by 0.035 (95% CI −0.006 to 0.076) from the raw score to V1, so we have no evidence of a drop. For GenAge, adjustment moved the AUROC from 0.44 to 0.51. In GenAge without yeast on genes measured in at least two lines (*n* = 139/87), V1, V6 and V7 gave AUROCs of 0.57–0.60, none significant after correction (q = 0.082 to 0.27 within the family). The GenAge mouse subset had 23/66 genes. Restricting CellAge to records based on loss of function (*n* = 129/343) gave 0.54 for R_rob_ (q = 0.27), 0.60 for the raw score (q = 0.0070) and 0.60 for proliferation alone (q = 0.0044). For the GTEx-derived signature, the multi-line GenAge test had *n* = 197/88, the all-gene GenAge test gave q = 0.27, and the multi-line CellAge test gave q = 0.050. Among GenAge orthologs, the odds ratio for DepMap common essentials was 1.37.

#### Multi-objective ranking

Pareto optimality used the five objectives in the protocol (R_rob_, sign consistency, identity preservation, absence of proliferation gain and absence of stress induction) on the 2,393 genes measured in at least two lines. The exploratory filter required on-target z < −0.5, response above the non-targeting 95th percentile in at least half of the measured screens, and at least 25 cells (mean over screens). The z < −0.5 threshold marks clear knockdown, given a median on-target z of about −0.7. The 25-cell threshold is a stricter quality filter than the 20 cells needed to form a pseudobulk profile. After this filter, 1,013 of the 2,393 genes remained. Before filtering, the 6 Pareto-optimal knockdowns with weak or unmeasured on-target knockdown included olfactory receptor genes that are not expressed in these lines. Under V6 the front had 17 genes, 11 of them shared with V1; under V7 it had 15, all shared. Of the 2,393 genes, 1,190 had R_rob_ > 0.

### QUANTIFICATION AND STATISTICAL ANALYSIS

H1a: one-sided Spearman tests between each pair of lines, using the K562 essential-gene screen for K562 (Holm^59^). H1b: within-line minus mean between-line ρ on the 1,784 knockdowns measured in all screens (2,000 bootstraps over knockdowns, with ranks computed once before resampling). H2: Spearman ρ of R with π (Holm). H3: paired one-sided Wilcoxon tests on absolute errors (Holm over screens; supported if significant in more than half). H4: one-sided Mann–Whitney tests of R_rob_ (Holm over two tests). H5a: 200 donor-level bootstraps, re-deriving the signature and all scores each time, with the median ρ against the full-cohort ranking compared with 0.8. H5b: one-sided Spearman test of agreement with the ranking from the GTEx-derived signature. Holm families were H1a (six pairs), H2 (five screens), H3a and H3b (five screens each) and H4 (two tests); we did not correct across hypotheses.

Exploratory analyses: 2,000 donor bootstraps for H5 and for every cohort specification; 500 degree-preserving randomized networks for H3a, evaluated on the same folds, with empirical *P* = (1 + number of nulls with ρ ≥ observed)/(1 + 500), and H3a re-tested against the mean prediction over all randomizations (Holm over five screens); paired DeLong tests and a logistic model (label ∼ proliferation shift + raw score) for AUROC comparisons; minimum detectable AUROC at 80% power; random signatures as above. Figure 5 and Figure S9 show Hanley–McNeil intervals. Bootstrap intervals (2,000 resamples over genes) are given in the text for the pre-specified tests. Network CIs used 2,000 bootstrap resamples of held-out knockdowns, and age-prediction CIs the Fisher z transformation. Table S5 lists all 410 exploratory tests with a P value in 17 families, each test once, with its P value and the Holm and Benjamini–Hochberg q values within its family; the top-k sweep forms its own family. The top-k sweep repeats the F0 test at other k; its q values describe the sweep and are not used to claim significance. Exploratory statistics without a P value (planted effects, split halves, reliability) are in Table S3. For each family we give Benjamini–Hochberg q values. In the text, an exploratory result is called significant only if q < 0.05 within its family; other exploratory P values are given as uncorrected. The extended run of 2,000 donor bootstraps gave a median ρ of 0.778 (95% of replicates 0.62 to 0.87) on all genes, and 0.856 (0.70 to 0.93; 81% of bootstraps at or above 0.8) on the genes measured in at least two lines. For H1b, the one-sided 95% bound of the difference was −0.04. Response size was far more reproducible between the two K562 screens (ρ = 0.88) than the score. The like-for-like ranking from discovery donors aged 20 to 79 correlated with the GTEx-fibroblast ranking at ρ = 0.01.

Tests on knockdown scores were rank-based. Per-gene age effects used ordinary least squares t-statistics, which assume roughly normal residuals. Rank ties received average ranks, and knockdowns with missing values were excluded from that analysis only. Bootstrap CIs over knockdowns treat knockdowns as independent, although related genes respond similarly, so these CIs may be too narrow; the block bootstrap addresses this for the network comparison only. Exact n, what n counts (donors, knockdowns or genes) and the CIs are given in the figure legends and in Tables S4, S14, S15 and S16.

#### Software, computation and reproducibility

Analyses used Python 3.13.7 with the package versions in the key resources table, on one workstation (8 CPU cores, 8 GB RAM, no GPU). Analysis code was written with help from Claude (Opus 5.5; Anthropic) and checked by unit tests on synthetic data, which cover the score and its target exclusion, residualization, Holm correction against statsmodels, the Pareto front, degree preservation of the network null, the robust-score definition, recovery of a planted age effect under covariates, and the generic two-table cohort loader; the tests run on every commit. Seeds and resample counts are fixed in one configuration file. Every figure panel has a source-data file, and the result files behind every reported number are in the Zenodo deposit. To run TRACE, provide an aging cohort as a genes-by-samples expression table with a sample table of age and sex, and a pseudobulk z-shift matrix per perturbation for each screen. The command python run_all.py runs all checks and writes one report table (results/trace_report.csv) with each check’s value, threshold and result (see the repository documentation).

## Supplemental information

**Figure S1.**
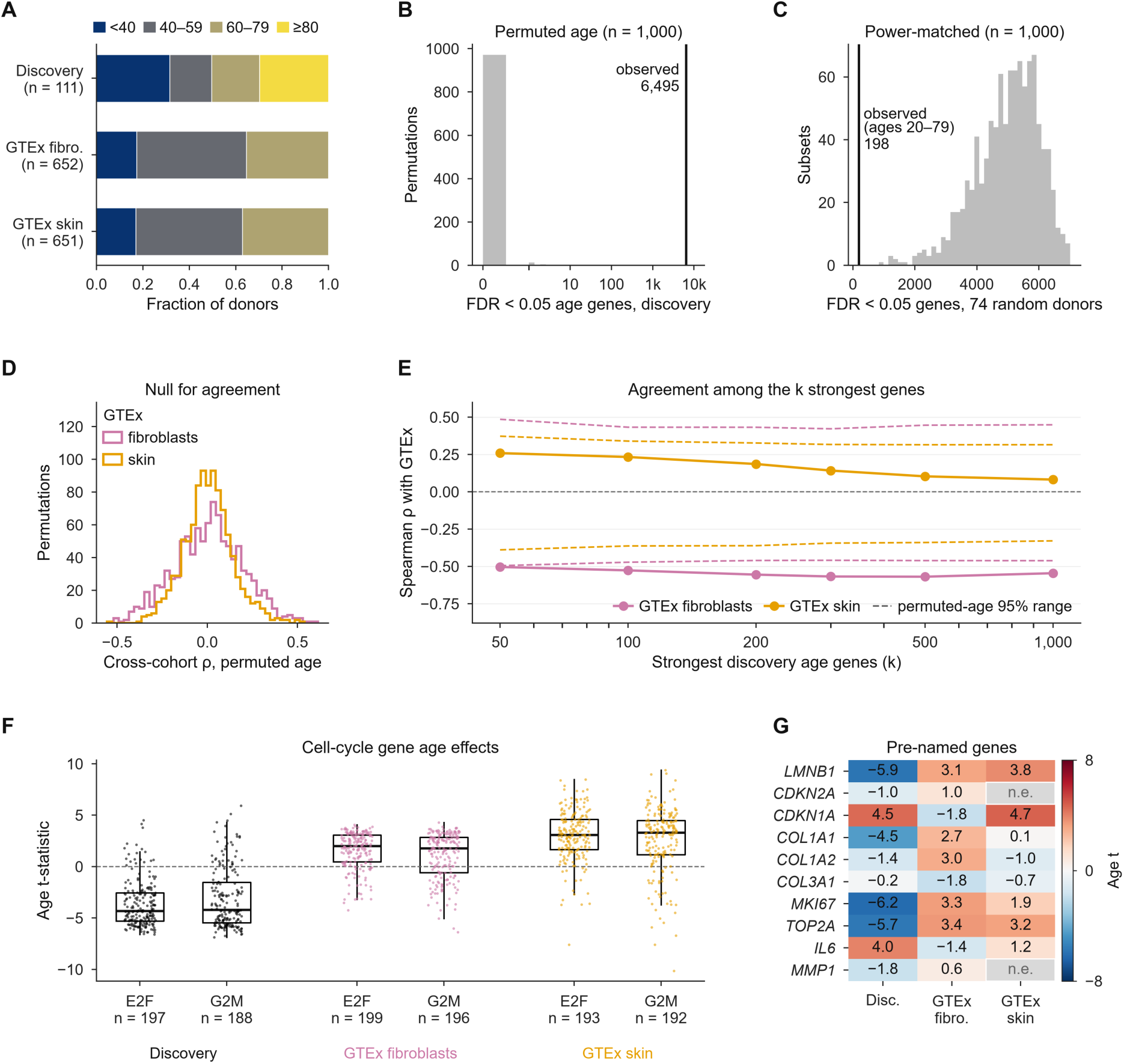
Cohorts and controls for the cohort comparison, related to Figure 1. (A) Donor age by cohort (*n* = 111, 652 and 651 donors); GTEx has no donors aged 80 or older. (B) Number of FDR < 0.05 age genes in the discovery cohort under 1,000 permuted age labelings (observed, 6,495; median 0, 95th percentile 0, 99th percentile 4; x axis on a symmetric log scale). (C) FDR < 0.05 age genes in 1,000 random subsets of 74 discovery donors (observed for donors aged 20–79, 198). (D) Genome-wide Spearman ρ of age t-statistics between the discovery cohort and GTEx fibroblasts or skin under 1,000 permuted age labelings (main model, F0 × G0). (E) Spearman ρ among the k strongest discovery age genes (k = 50 to 1,000) with GTEx fibroblasts (pink) and skin (orange), main model; dashed lines, 95% range under 1,000 permuted age labelings at each k (the same permutations as the F0 test in Table S4); values in Table S3. (F) Age t-statistics of Hallmark E2F-target and G2M-checkpoint genes in each cohort (box, quartiles; whiskers, 1.5 × interquartile range; points, genes; n, genes). (G) Age t-statistics of genes named before the analysis (n.e., not expressed); color, age t; values in Table S6. All panels except (A) are exploratory.

**Figure S2.**
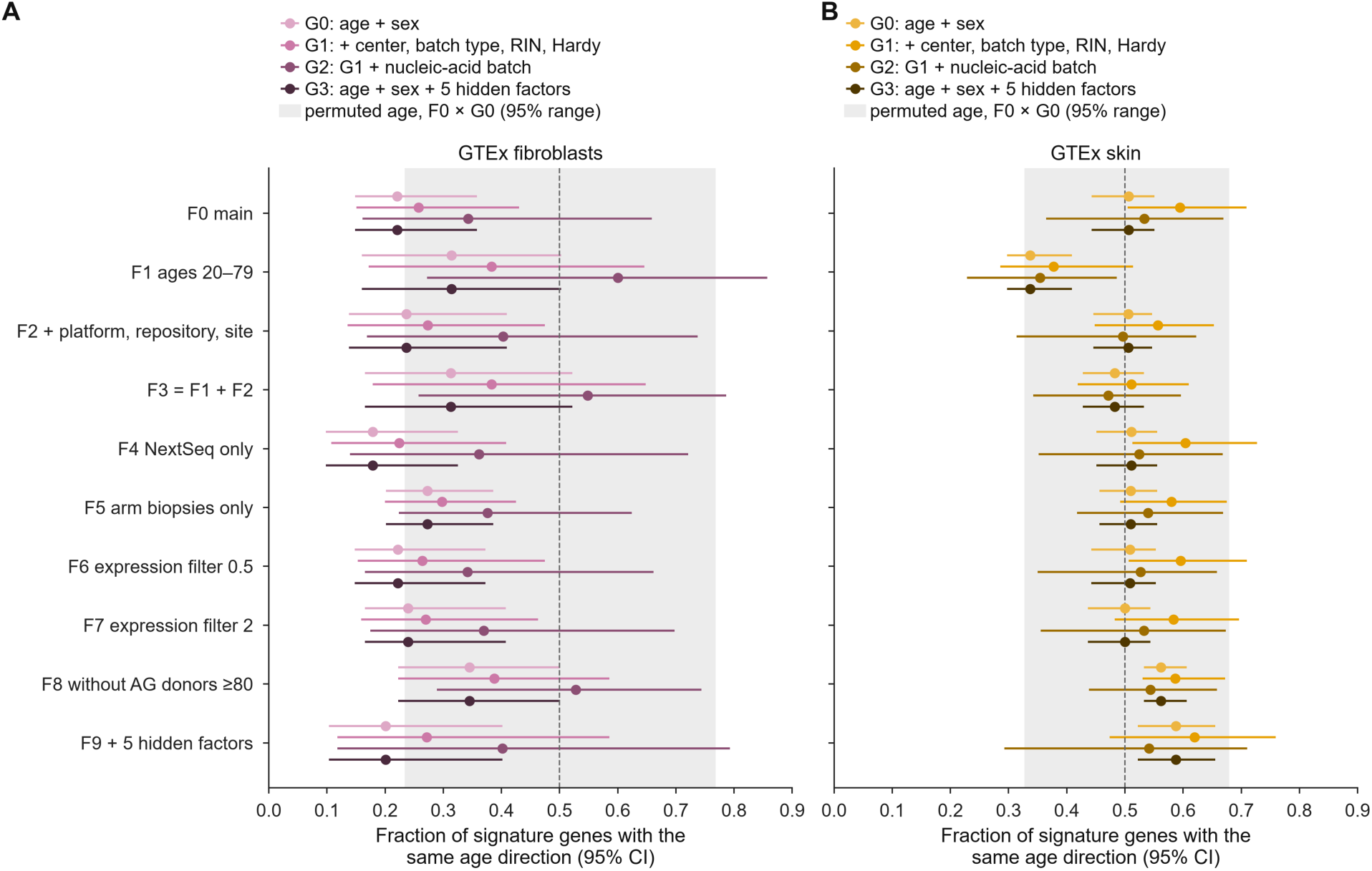
Sign agreement of signature genes across cohort specifications, related to Figure 1. Fraction of discovery signature genes with the same age direction in GTEx fibroblasts (A) and skin (B) for every discovery specification (F0–F9, rows) and replication specification (G0–G3, shades); error bars, 95% bias-corrected CIs from 2,000 donor bootstraps; dashed line, 0.5; gray band, 95% range of sign agreement for F0 × G0 under 1,000 permuted age labelings (0.23 to 0.77 for fibroblasts and 0.33 to 0.68 for skin). G3 sign agreement equals G0 by construction: the five G3 hidden factors are computed from residuals after the age + sex model, so they are orthogonal to age and leave the sign of every gene’s age effect unchanged; they change only the standard errors, and so the G3 ρ values. F0 × G0 is the main model. AG, NIA Aging Cell Repository. Exploratory; values in Table S4 and Table S3.

**Figure S3.**
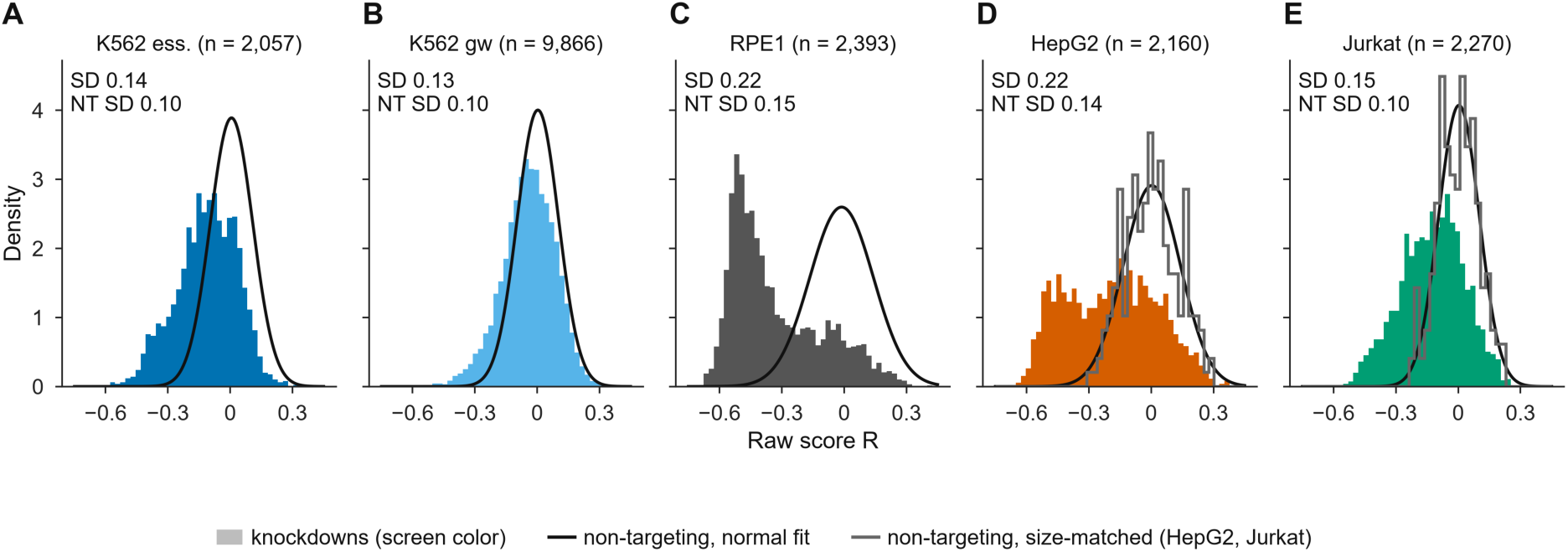
Score distributions, related to Figure 2. (A–E) Histograms of the raw score R for knockdowns in each screen (colors, screens; n, knockdowns); black line, normal fit to non-targeting entries; gray line (HepG2, Jurkat), non-targeting groups size-matched to a typical knockdown. The SD of knockdowns and of non-targeting entries is given above each panel. In RPE1 and HepG2 the whole knockdown distribution is shifted to about −0.5. Exploratory.

**Figure S4.**
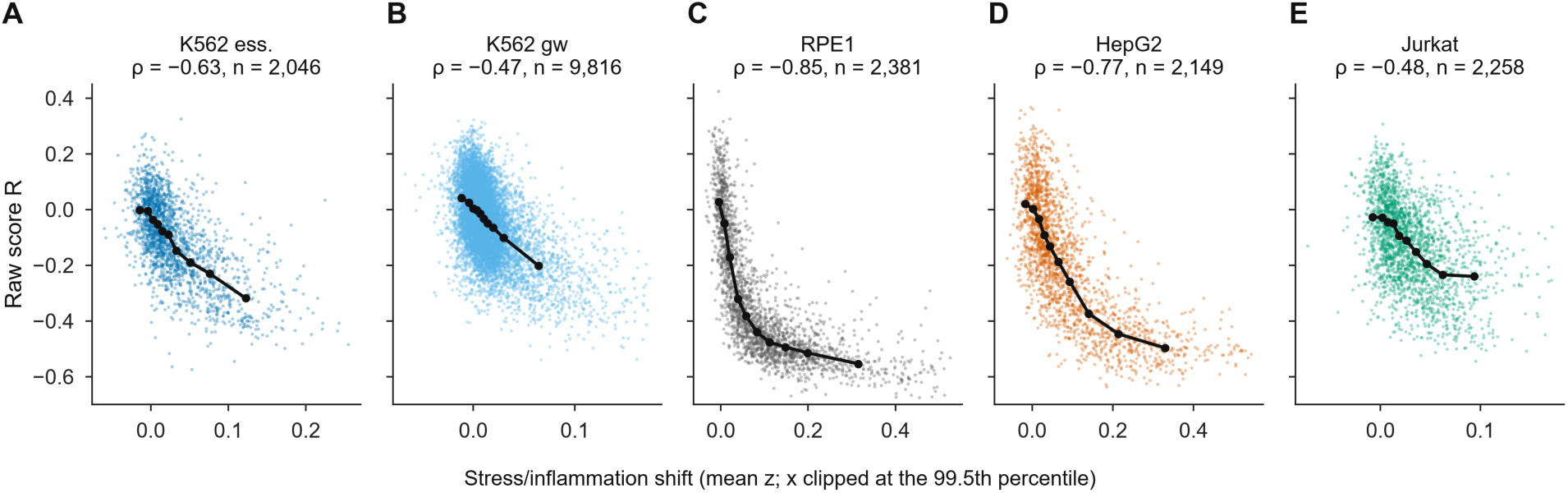
Raw scores and the stress program, related to Figure 2. (A–E) Raw score R versus stress/inflammation shift (mean z over Hallmark TNF-α signaling via NF-κB, inflammatory response and p53 pathway genes), one point per knockdown; black line, decile medians; ρ, Spearman correlation; n, knockdowns shown (x axis clipped at the 99.5th percentile). Exploratory.

**Figure S5.**
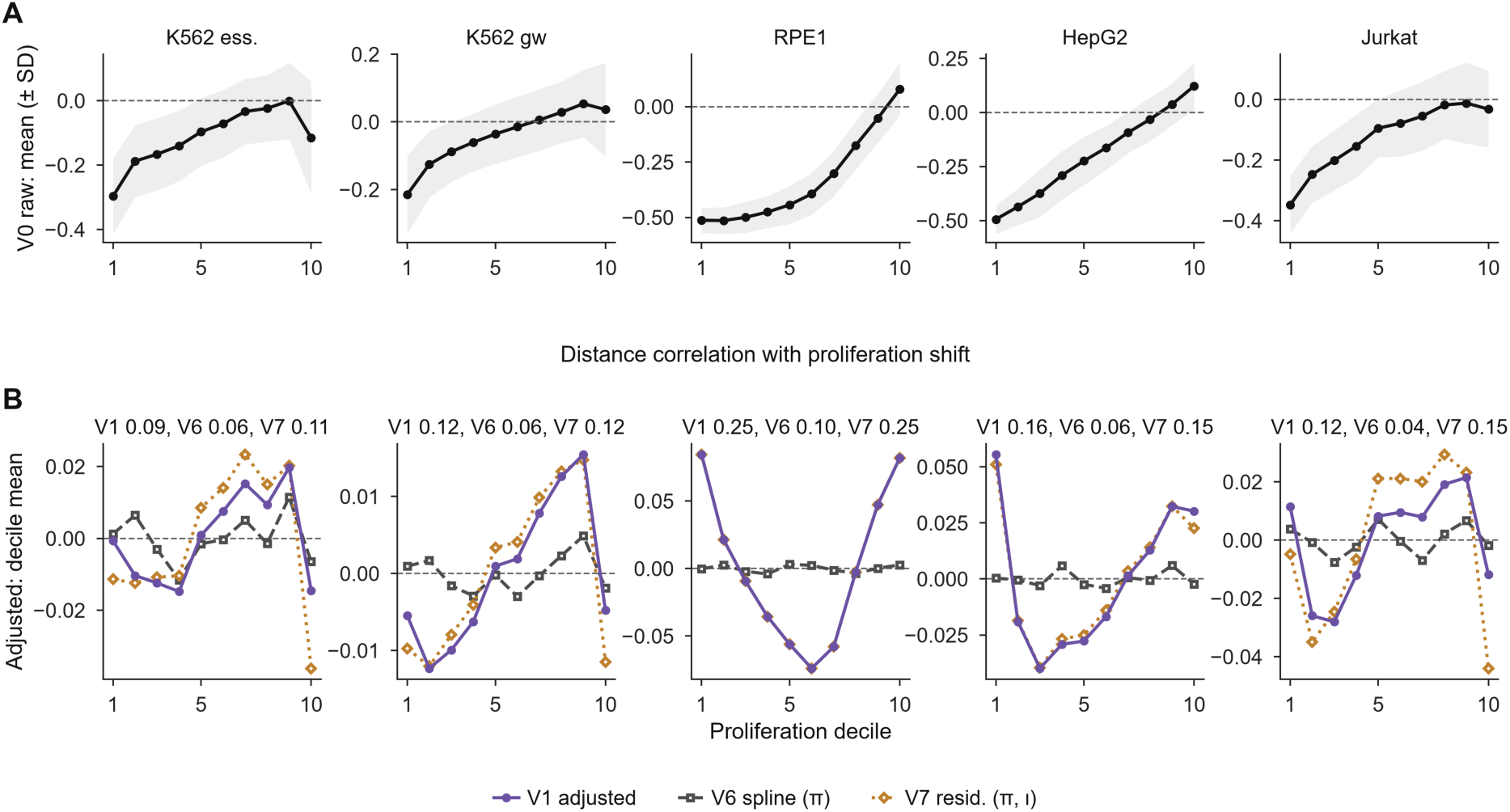
Proliferation dependence by decile, related to Figure 2. (A) Mean of the raw score (V0) by decile of proliferation shift in each screen; gray band, ± 1 SD. (B) Decile means of V1, V6 and V7, with the distance correlation of each definition with proliferation shift above each panel; V1 and V7 overlap in RPE1 and HepG2. Exploratory; values for every definition, with mutual information, in Table S9.

**Figure S6.**
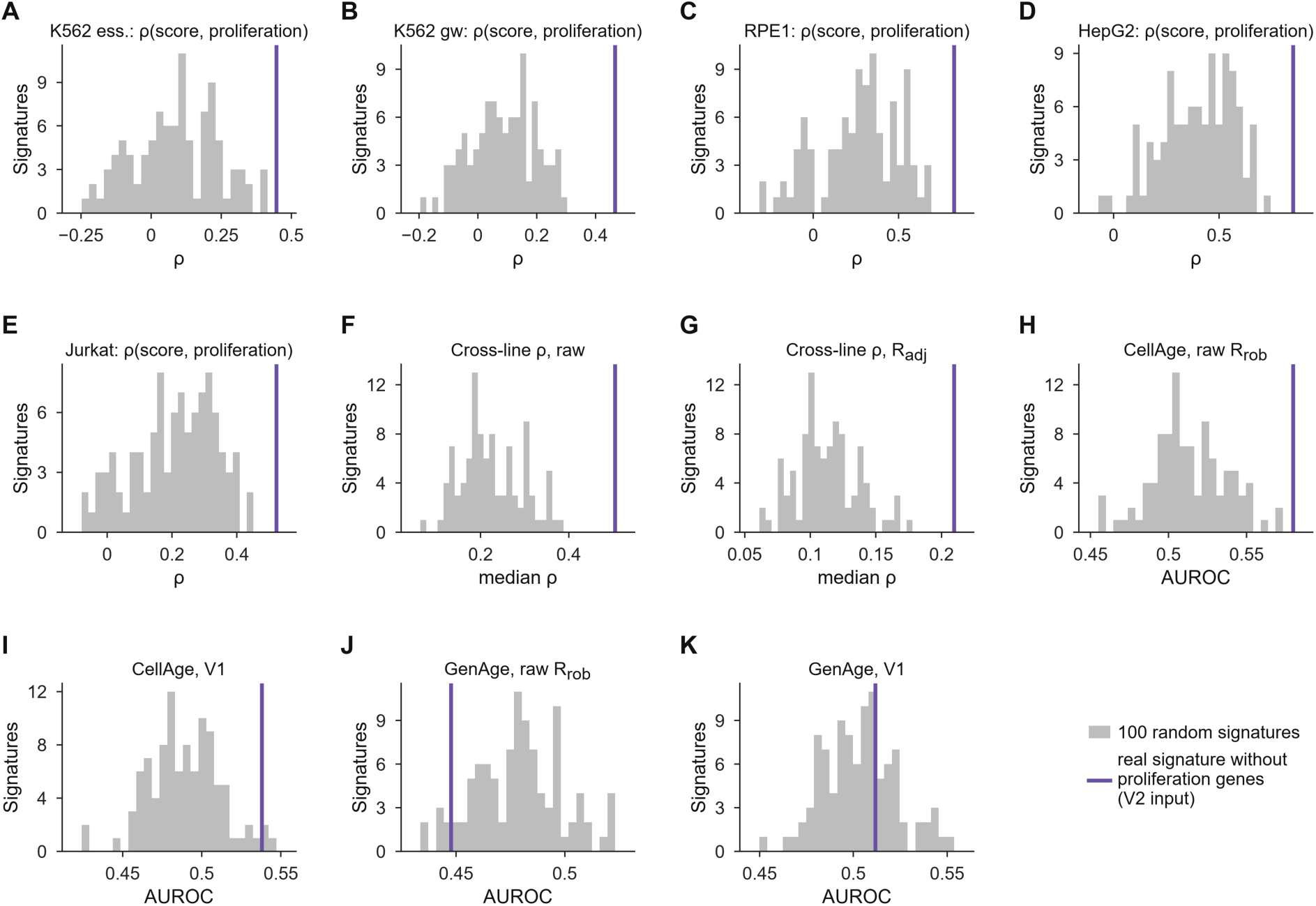
Random-signature nulls, related to Figure 2. Distributions over 100 random expression- and variance-matched signatures drawn from genes outside the proliferation sets (bars); vertical line, the real signature without its proliferation genes (V2 input), so that neither side contains cell-cycle genes. Because the real values use the V2 input, they differ from the raw-score values in the Results (for example, raw CellAge AUROC 0.58 here and 0.57 with the full signature). (A–E) ρ of the score with proliferation per screen. (F and G) Median cross-line ρ for the raw score and for R_adj_. (H–K) AUROC for CellAge and GenAge labels with the raw score and with V1. The real signature exceeds all random signatures in (A–H) (one-sided empirical *P* = 0.0099, the smallest attainable with 100 draws; q = 0.014 within the family, Table S5) and 97% of them in (I); for GenAge it does not (J and K). Values in Table S3. Exploratory.

**Figure S7.**
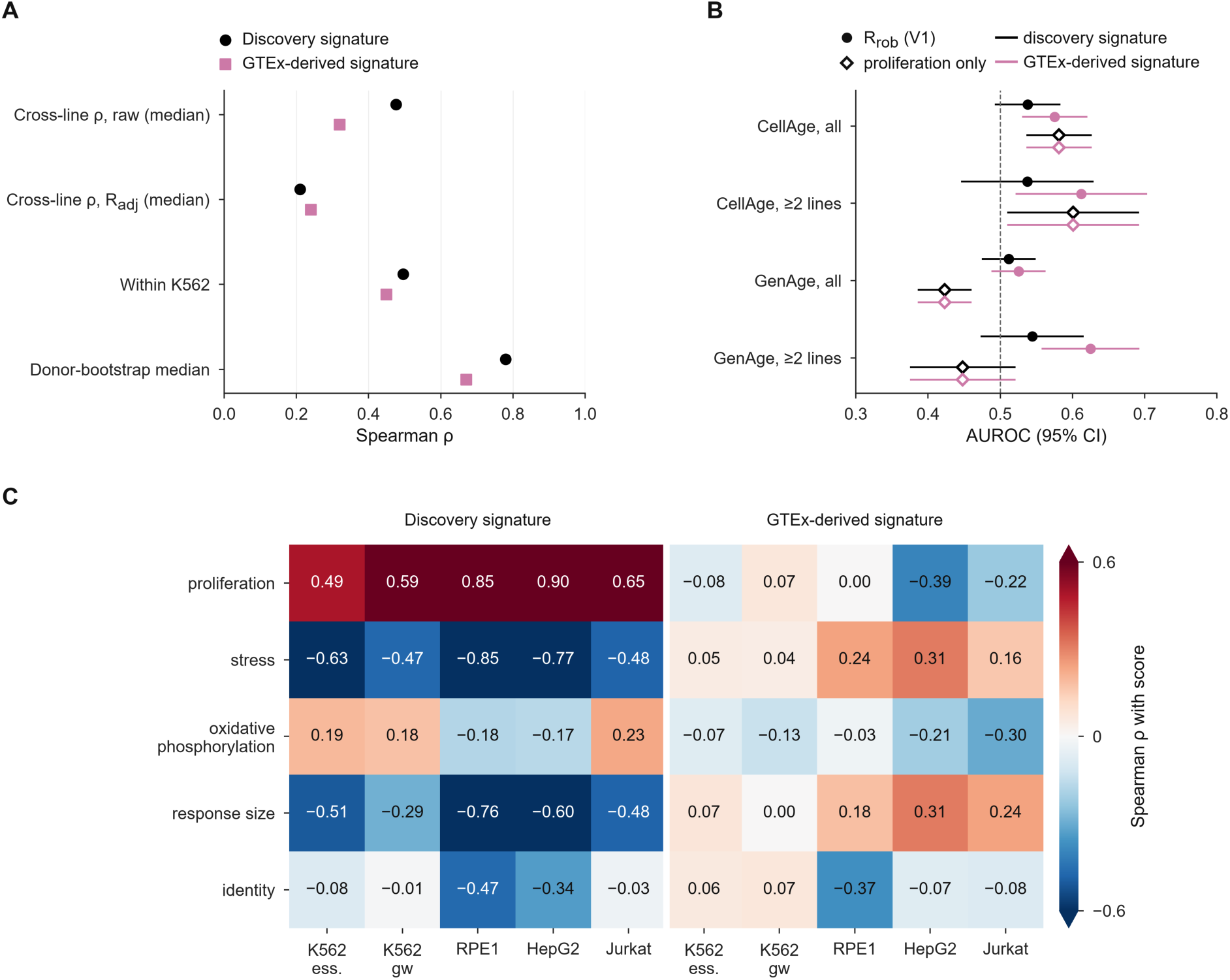
TRACE run with a GTEx-derived signature, related to Figure 2. (A) Agreement statistics for the discovery signature (circles) and the GTEx-derived signature, built the same way from GTEx fibroblasts (squares): median cross-line ρ, within-K562 ρ and the donor-bootstrap median. (B) AUROC (95% Hanley–McNeil CI) for CellAge and GenAge labels on all genes and on genes measured in at least two lines, for R_rob_ (V1; filled) and proliferation shift alone (open); dashed line, chance. (C) Spearman ρ of each signature’s raw score with broad knockdown programs per screen. Exploratory; values in Table S11.

**Figure S8.**
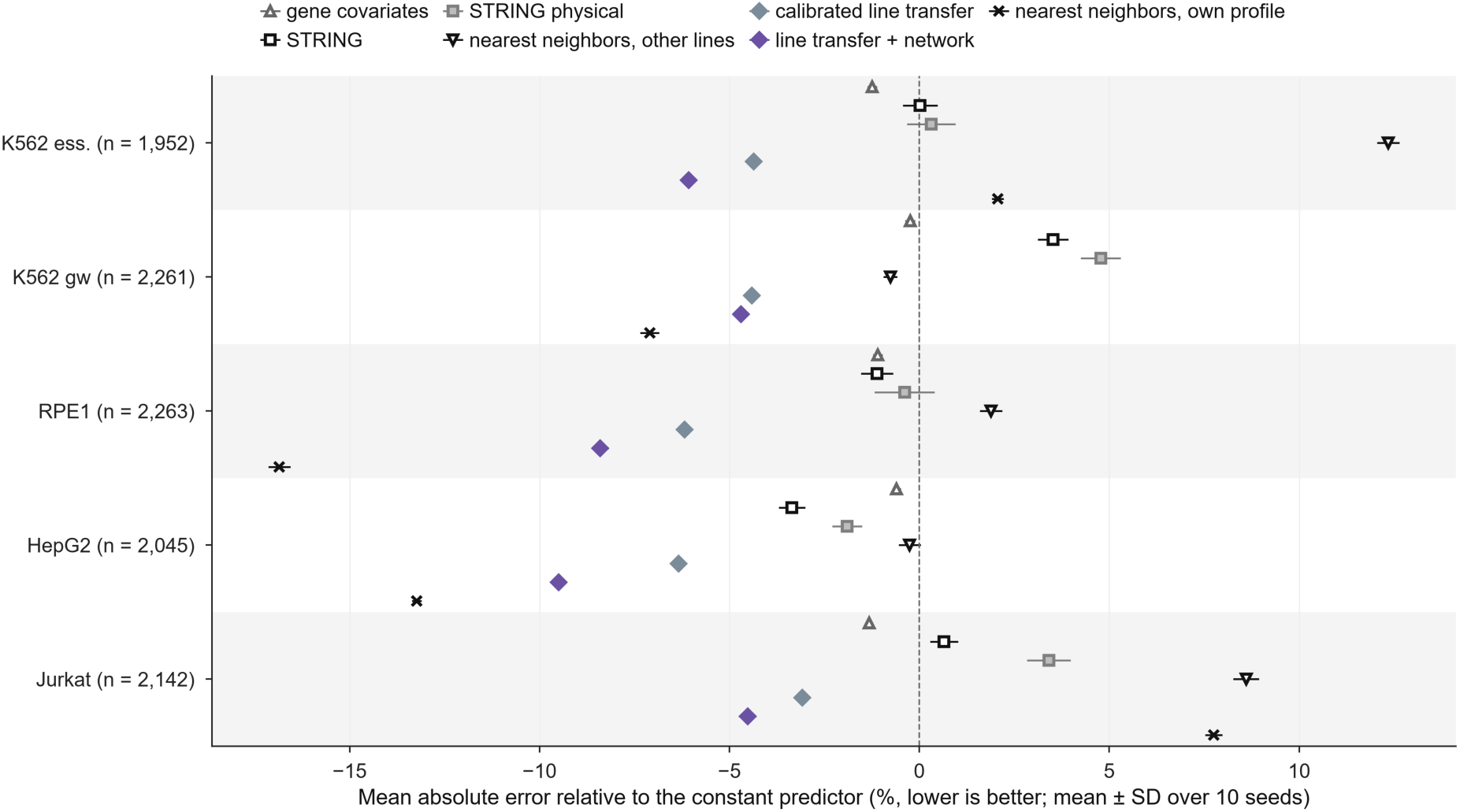
Prediction error of all network baselines, related to Figure 4. Mean absolute error of each predictor relative to the constant predictor (training mean), per screen, on the common subset (n, knockdowns); points, mean over 10 cross-validation seeds; error bars, SD. The nearest-neighbor predictor that uses the knockdown’s own profile is a reference, not a competitor for unprofiled genes. Exploratory; values in Table S14.

**Figure S9.**
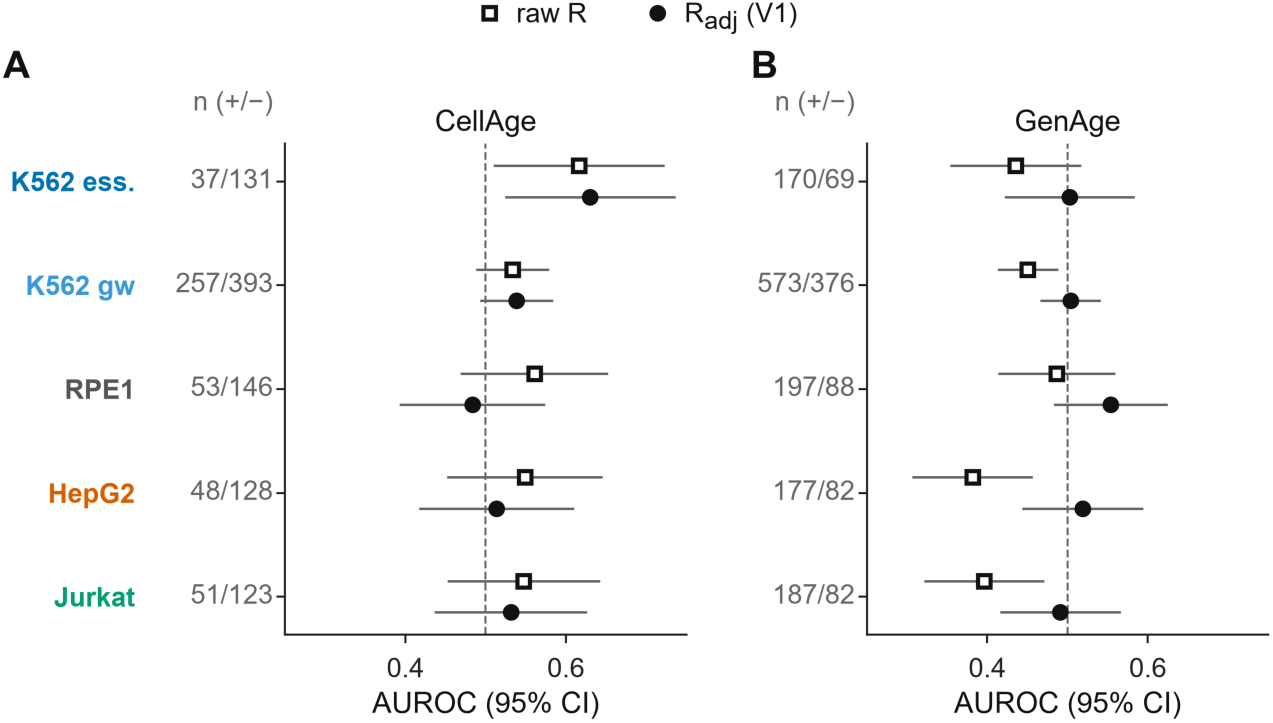
Single-screen evidence tests, related to Figure 5. AUROC (95% Hanley–McNeil CI) of the raw score (open squares) and the per-screen adjusted score R_adj_ (V1; filled circles) for CellAge (A) and GenAge (B) labels in each screen; *n* (+/−), positive/negative genes. Counts for K562 genome-wide almost equal the pooled counts. The only adjusted CI above 0.5 is CellAge in K562 ess. (0.63 (95% CI 0.52 to 0.74); *n* = 37/131), which does not survive correction over the 20 tests in this figure (Table S5). Exploratory.

**Figure S10.**
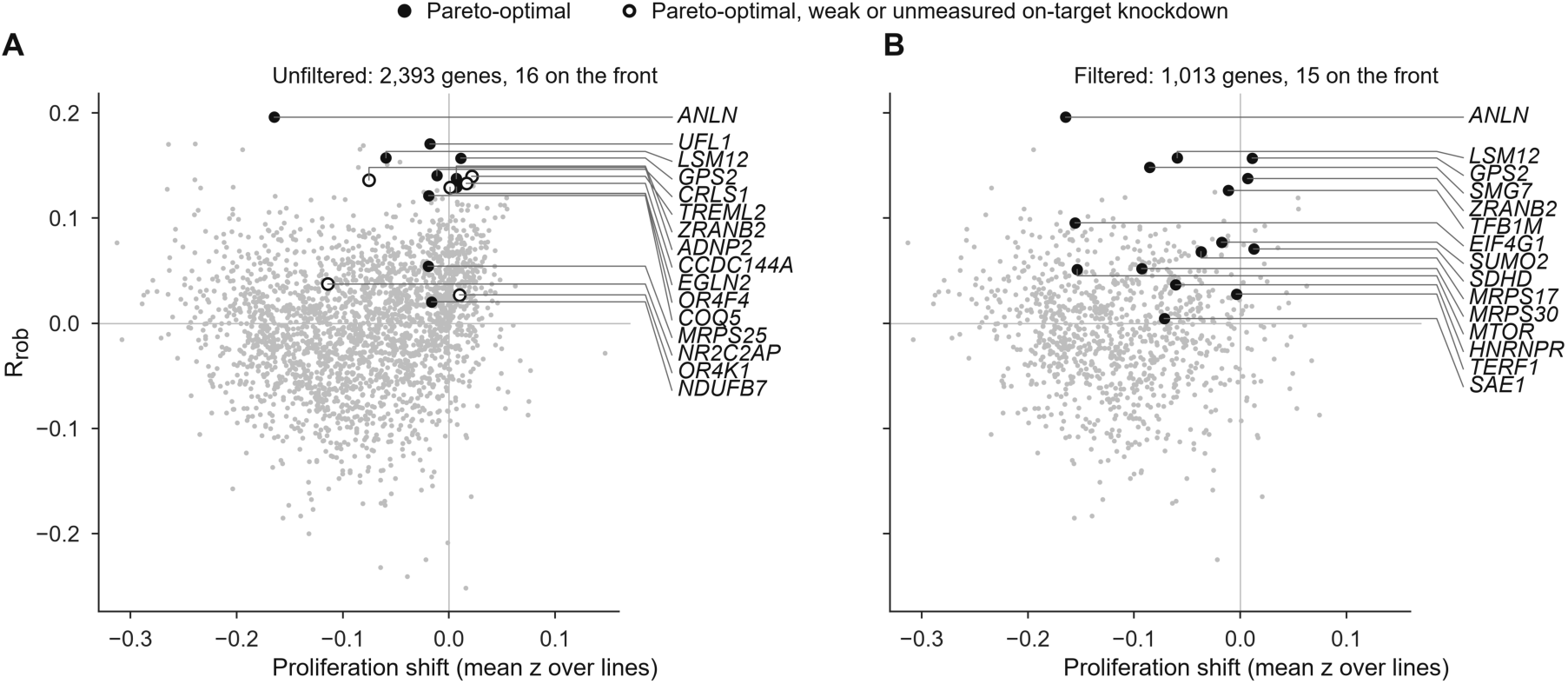
Pareto front before and after filtering, related to STAR Methods. R_rob_ versus proliferation shift for the 2,393 genes measured in at least two lines. (A) Unfiltered: 16 genes on the front (filled; open, weak or unmeasured on-target knockdown). (B) After filtering on on-target z < −0.5, response above the non-targeting 95th percentile in at least half of the measured screens and at least 25 cells: 1,013 genes, 15 on the front. Pareto status uses five objectives, two of which are shown. Every front gene is named, ordered by R_rob_, with a leader line to its point. *ANLN*, a CellAge senescence inhibitor whose knockdown slows growth, leads the unfiltered front. Exploratory; values in Table S18.

**Table S1.** Positioning of TRACE relative to the closest prior work, related to Table 1. Entries describe each study’s main aim and data as we understand them; they are not claims about what those studies did not test.

| Study | Question | Aging reference | Perturbation data | Main output | What TRACE adds for the same data type |
|---|---|---|---|---|---|
| Janssens et al. 2019<sup>7</sup> | Which compounds oppose a human transcriptomic age signature? | Transcriptomic age signature | Connectivity Map profiles | Candidate compounds; HSP90 inhibition tested in *C. elegans* | Cross-cohort signature test; proliferation-adjusted scores |
| Tyshkovskiy et al. 2019, 2023<sup>8,9</sup> | Shared signatures of lifespan-extending interventions | Longevity-intervention signatures | Connectivity Map (mimetics) | Longevity signatures; candidate mimetics | Genetic knockdowns across lines; replicate and cohort tests |
| REVIVE (Jung et al. 2025)<sup>10</sup> | Which perturbations restore a youthful transcriptional state? | Transcriptional aging clock; young-vs-old differentially expressed genes | >10,000 chemical and genetic perturbations | Ranked perturbations; aging-hallmark enrichment | Tests fixed in advance; cohort swap; proliferation-only label baseline |
| Tadevosyan et al. 2025<sup>11</sup> | Can a foundation model nominate anti-aging gene perturbations? | scGPT fine-tuned on single-cell age groups | In silico perturbation | Candidate pro-/anti-aging genes | Measured knockdown responses; network and transfer baselines |
| Perturbation-model benchmarks<sup>29</sup> | Do deep models beat simple baselines? | Not aging-specific | Perturb-seq | Baselines are hard to beat | Aging end point; network vs randomized network and transfer |

**Table S2.**
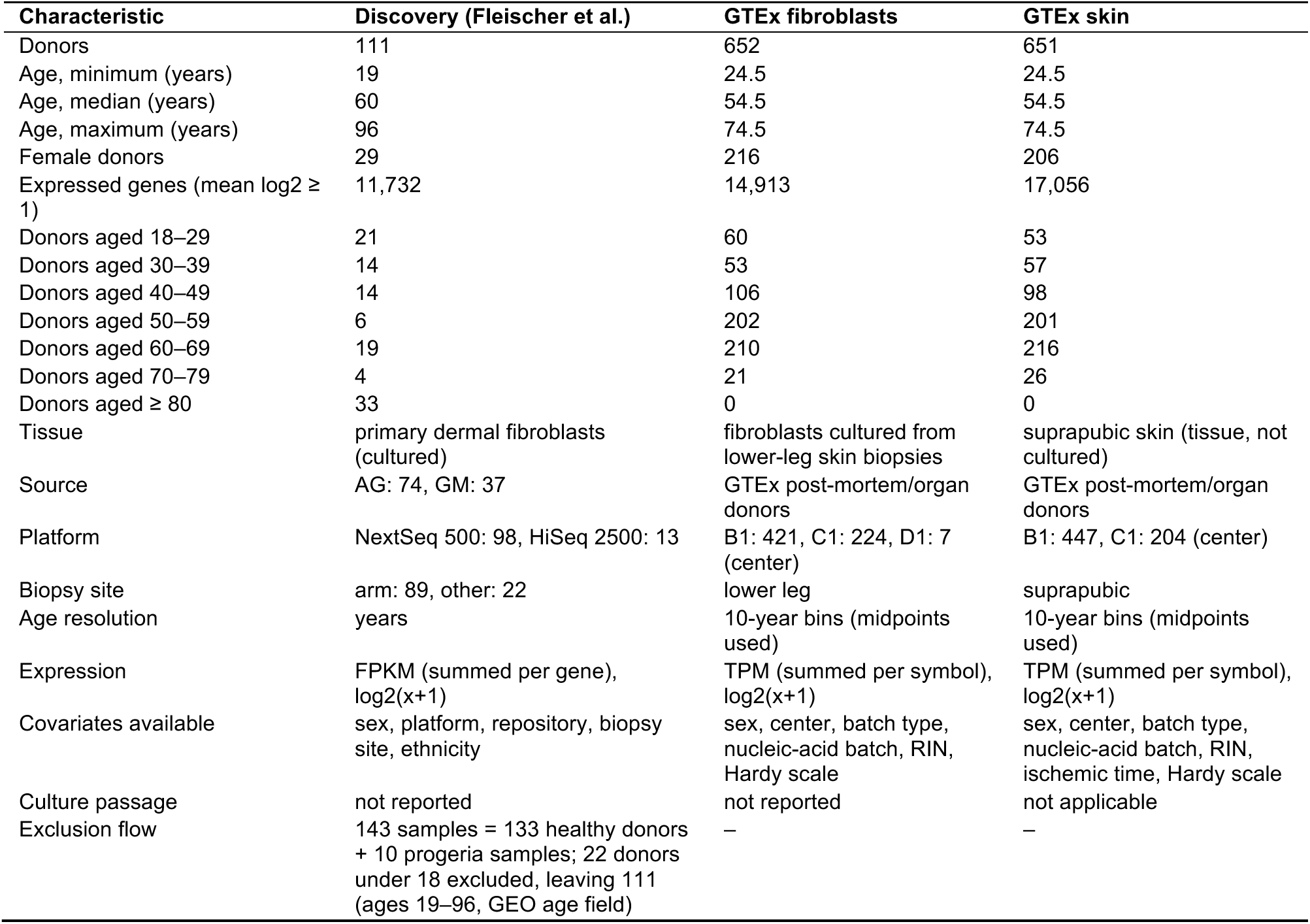
Aging cohorts, related to Figure 1. Source and platform entries are numbers of donors. AG, NIA Aging Cell Repository; GM, NIGMS Human Genetic Cell Repository; GTEx centers B1, C1 and D1 as coded by GTEx.

**Table S6.**
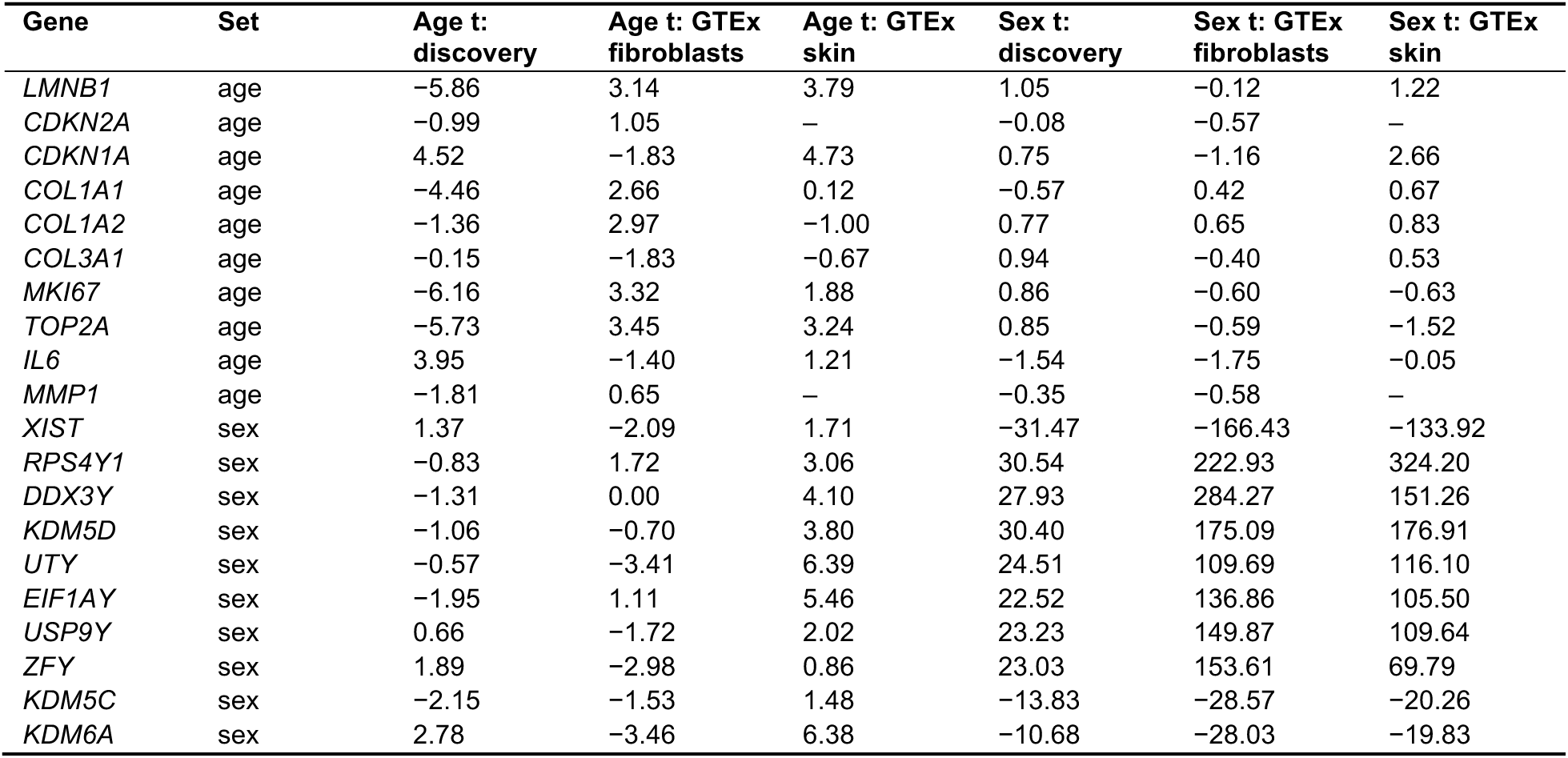
Age and sex t-statistics of genes named before the analysis, related to Figure 1. t, ordinary least squares t-statistic (main model); –, not expressed. Exploratory.

**Table S7.** Sex positive control with permuted-age and permuted-sex nulls, related to Figure 1. Main model (F0 × G0); 2,000 donor bootstraps; 1,000 permuted age labelings for age and 1,000 permuted sex labelings for sex; two-sided empirical P, uncorrected. Exploratory.

| **GTEx cohort** | **Effect and gene set** | **Spearman ρ** | **95% CI (bias-corrected)** | **Null, 95% range** | **Permutation *P*** |
|---|---|---|---|---|---|
| GTEx fibroblasts | Age, all genes | −0.28 | −0.46 to −0.09 | −0.40 to 0.39 | 0.16 |
| GTEx fibroblasts | Age, 200 strongest genes | −0.55 | −0.72 to −0.47 | −0.46 to 0.43 | 0.011 |
| GTEx fibroblasts | Sex, all genes | 0.19 | 0.07 to 0.44 | −0.38 to 0.37 | 0.31 |
| GTEx fibroblasts | Sex, 200 strongest genes | 0.65 | 0.52 to 0.80 | −0.46 to 0.46 | 0.002 |
| GTEx fibroblasts | Sex, 200 strongest autosomal genes | 0.34 | 0.11 to 0.63 | −0.46 to 0.45 | 0.14 |
| GTEx fibroblasts | Sex, all autosomal genes | 0.17 | – | – | – |
| GTEx skin | Age, all genes | 0.15 | 0.03 to 0.29 | −0.28 to 0.28 | 0.28 |
| GTEx skin | Age, 200 strongest genes | 0.19 | 0.02 to 0.51 | −0.36 to 0.33 | 0.25 |
| GTEx skin | Sex, all genes | −0.12 | −0.31 to 0.06 | −0.26 to 0.25 | 0.37 |
| GTEx skin | Sex, 200 strongest genes | 0.32 | 0.17 to 0.48 | −0.32 to 0.34 | 0.057 |
| GTEx skin | Sex, 200 strongest autosomal genes | −0.18 | −0.44 to 0.02 | −0.31 to 0.34 | 0.25 |
| GTEx skin | Sex, all autosomal genes | −0.13 | – | – | – |

**Table S8.** Cross-cohort elastic-net age prediction, related to Figure 1. Elastic net (l1 ratio 0.5) trained on one cohort; genes used = genes with a non-zero coefficient. MAE, mean absolute error. Exploratory.

| Training cohort | Test cohort | *n* (test donors) | MAE (years) | Pearson r | Genes used |
|---|---|---|---|---|---|
| Discovery | Discovery (five-fold cross-validation) | 111 | 13.7 | 0.72 | 358 |
| Discovery | GTEx fibroblasts | 652 | 20.9 | −0.01 | 358 |
| Discovery | GTEx skin | 651 | 13.2 | 0.06 | 358 |
| GTEx fibroblasts | GTEx fibroblasts (five-fold cross-validation) | 652 | 8.8 | 0.54 | 340 |
| GTEx fibroblasts | Discovery | 111 | 20.6 | 0.29 | 340 |
| GTEx fibroblasts | GTEx skin | 651 | 10.8 | 0.24 | 340 |
| GTEx skin | GTEx skin (five-fold cross-validation) | 651 | 8.2 | 0.63 | 585 |
| GTEx skin | Discovery | 111 | 22.8 | −0.09 | 585 |
| GTEx skin | GTEx fibroblasts | 652 | 11.0 | 0.17 | 585 |

**Table S10.**
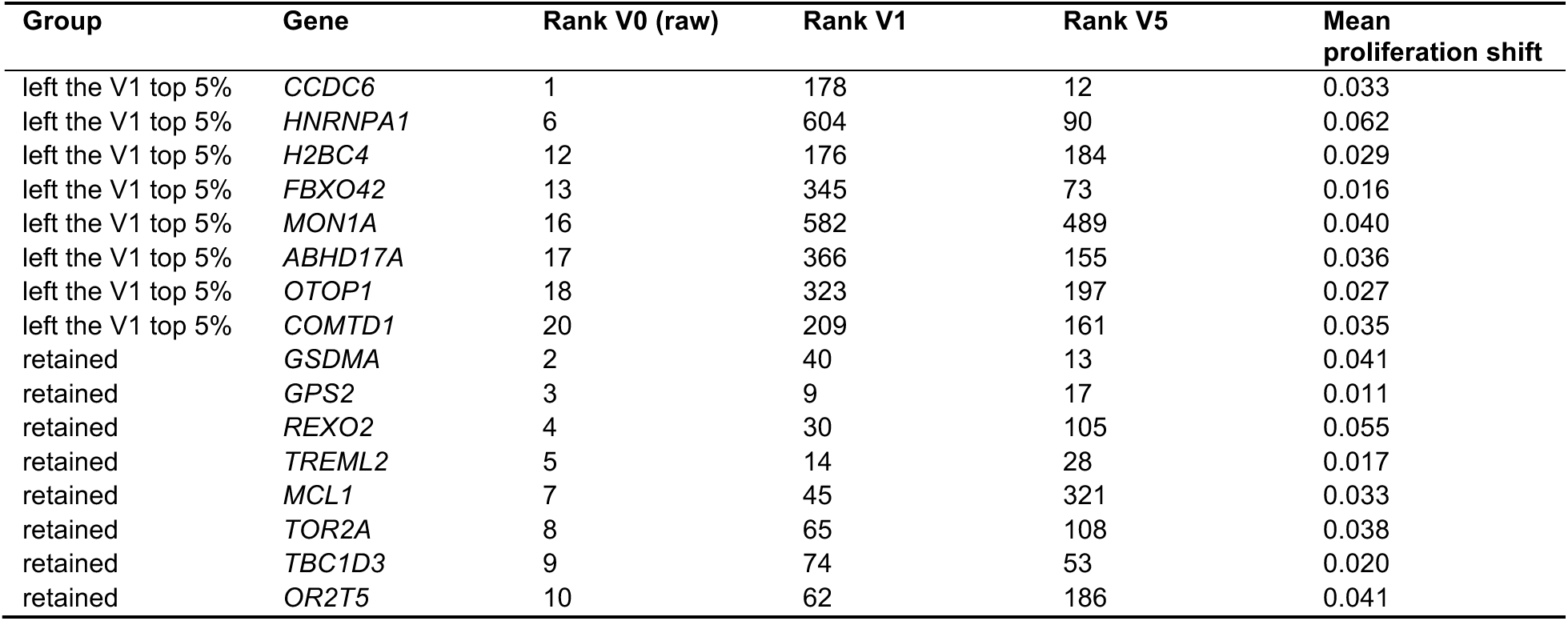
Raw top candidates that leave or stay in the V1 top 5%, related to Figure 2. k = 119 of the 2,393 genes measured in at least two lines; ranks of R_rob_ under each definition; *H2BC4* was HIST1H2BC. Neither group has been tested experimentally. Exploratory.

| Group | Gene | Rank V0 (raw) | Rank V1 | Rank V5 | Mean proliferation shift |
|---|---|---|---|---|---|
| left the V1 top 5% | *CCDC6* | 1 | 178 | 12 | 0.033 |
| left the V1 top 5% | *HNRNPA1* | 6 | 604 | 90 | 0.062 |
| left the V1 top 5% | *H2BC4* | 12 | 176 | 184 | 0.029 |
| left the V1 top 5% | *FBXO42* | 13 | 345 | 73 | 0.016 |
| left the V1 top 5% | *MON1A* | 16 | 582 | 489 | 0.040 |
| left the V1 top 5% | *ABHD17A* | 17 | 366 | 155 | 0.036 |
| left the V1 top 5% | *OTOP1* | 18 | 323 | 197 | 0.027 |
| left the V1 top 5% | *COMTD1* | 20 | 209 | 161 | 0.035 |
| retained | *GSDMA* | 2 | 40 | 13 | 0.041 |
| retained | *GPS2* | 3 | 9 | 17 | 0.011 |
| retained | *REXO2* | 4 | 30 | 105 | 0.055 |
| retained | *TREML2* | 5 | 14 | 28 | 0.017 |
| retained | *MCL1* | 7 | 45 | 321 | 0.033 |
| retained | *TOR2A* | 8 | 65 | 108 | 0.038 |
| retained | *TBC1D3* | 9 | 74 | 53 | 0.020 |
| retained | *OR2T5* | 10 | 62 | 186 | 0.041 |

**Table S11.** Main tests and program links for the discovery and GTEx-derived signatures, related to Figure 2. The same code was run with each signature as the discovery input; for the GTEx-derived signature every test is exploratory. For the discovery signature, values in parentheses are Holm P within the two-test H4 family; for the GTEx-derived signature they are q values within its family in Table S5. Ranges over screens or pairs are given as low to high. The GTEx-derived signature cannot be compared with itself for H5b.

| **Test or statistic** | **Discovery signature** | **GTEx-derived signature** |
|---|---|---|
| H1a: raw cross-line ρ (range over pairs) | 0.42 to 0.64 | 0.22 to 0.41 |
| H1b: within-K562 versus between-line ρ | 0.50 versus 0.51 | 0.45 versus 0.33 |
| Adjusted cross-line ρ (range over pairs) | 0.15 to 0.26 | 0.18 to 0.35 |
| H2: ρ(raw score, proliferation), range over screens | 0.49 to 0.90 | −0.39 to 0.07 |
| H3a: network versus 10 nulls (screens significant) | 3/5 | 4/5 |
| H3b: line transfer + network versus line transfer (screens significant) | 5/5 | 5/5 |
| H4a: CellAge AUROC | 0.538 (Holm *P* = 0.10) | 0.575 (q = 0.0092) |
| H4b: GenAge AUROC | 0.512 (Holm *P* = 0.27) | 0.525 (q = 0.27) |
| H5a: donor-bootstrap median ρ (threshold 0.8) | 0.779 | 0.671 |
| H5b: ranking versus GTEx-derived signature | −0.05 (top 5% kept 4%) | – |
| Ranking versus GTEx skin signature (exploratory) | 0.19 (top 5% kept 11%) | −0.11 (top 5% kept 4%) |
| ρ(raw score, stress), range over screens | −0.85 to −0.47 | 0.04 to 0.31 |
| ρ(raw score, oxidative phosphorylation), range over screens | −0.18 to 0.23 | −0.30 to −0.03 |
| ρ(raw score, response size), range over screens | −0.76 to −0.29 | 0.00 to 0.31 |
| ρ(raw score, identity), range over screens | −0.47 to −0.01 | −0.37 to 0.07 |
| Cell-cycle genes in the signature (all; down arm) | 12.4%; 23.2% | 2.6%; 3.6% |

**Table S12.** Score reliability by screen and definition, related to Figure 3. Split cells: 20 random splits of each knockdown’s cells, Spearman–Brown corrected (mean). K562: Spearman ρ between the two K562 screens, which is also the within-K562 agreement. RPE1 not computed (pseudobulk release only). Exploratory.

| **Score definition** | **HepG2 (split cells)** | **Jurkat (split cells)** | **K562 (test–retest)** |
|---|---|---|---|
| V0 raw | 0.84 | 0.72 | 0.49 |
| V1 main | 0.64 | 0.62 | 0.38 |
| V2 genes removed | 0.85 | 0.76 | 0.51 |
| V3 resid. (all 3) | 0.64 | 0.59 | 0.34 |
| V4 resid. (prolif.) | 0.64 | 0.71 | 0.50 |
| V5 stratified | 0.56 | 0.65 | 0.45 |
| V6 spline resid. | 0.63 | 0.61 | 0.39 |
| V7 resid. (prolif. + identity) | 0.63 | 0.65 | 0.39 |

**Table S15.** Evidence labels and ortholog mapping, related to Figure 5. CellAge positive = senescence inducers, negative = inhibitors; GenAge positive = anti-longevity, negative = pro-longevity. For CellAge, counts are genes; for each GenAge organism, Positive and Negative are model-organism genes before ortholog mapping, so they differ from the human ortholog counts used in the tests (for example, mouse 45/85 organism genes map to 47/91 human orthologs; Table S16). For screens, the counts are positive/negative genes measured in that screen. Of the 911 yeast genes, 400 carry a longevity label (349 anti- and 51 pro-longevity; most others are annotated as necessary for fitness), and 246 of these map to human orthologs.

| Source | Subset | Records | Genes | Positive | Negative | Dropped (conflicting or ties) | Evaluable positive (mapped) | Evaluable negative (mapped) |
|---|---|---|---|---|---|---|---|---|
| CellAge | all records | 949 | 866 | 370 | 475 | 21 | 257 | 394 |
| GenAge | *Mus musculus* | 136 | 136 | 45 | 85 | – | – | – |
| GenAge | *Caenorhabditis elegans* | 889 | 887 | 578 | 283 | – | – | – |
| GenAge | *Drosophila melanogaster* | 202 | 201 | 73 | 125 | – | – | – |
| GenAge | *Saccharomyces cerevisiae* | 911 | 911 | 349 | 51 | – | – | – |
| GenAge | human consensus | – | 1,539 | 942 | 548 | 49 | 573 | 376 |
| Measured in screen | K562 ess. | – | 2,057 | CellAge 37/131 | GenAge 170/69 | – | – | – |
| Measured in screen | K562 gw | – | 9,866 | CellAge 257/393 | GenAge 573/376 | – | – | – |
| Measured in screen | RPE1 | – | 2,393 | CellAge 53/146 | GenAge 197/88 | – | – | – |
| Measured in screen | HepG2 | – | 2,160 | CellAge 48/128 | GenAge 177/82 | – | – | – |
| Measured in screen | Jurkat | – | 2,270 | CellAge 51/123 | GenAge 187/82 | – | – | – |

**Table S17.** Ranks of known senescence-gene knockdowns under each score definition, related to Figure 5. Rank 1 = highest score among the 9,870 ranked genes; –, not targeted in any screen.

| Gene | Lines measured | V0 | V1 | V2 | V3 | V4 | V5 | V6 | V7 | Genes ranked |
|---|---|---|---|---|---|---|---|---|---|---|
| *CDKN1A* | 0 | – | – | – | – | – | – | – | – | 9,870 |
| *CDKN2A* | 0 | – | – | – | – | – | – | – | – | 9,870 |
| *TP53* | 1 | 8,006 | 7,696 | 7,054 | 8,836 | 9,235 | 9,057 | 7,731 | 7,373 | 9,870 |
| *RB1* | 1 | 128 | 1,937 | 404 | 764 | 548 | 455 | 1,757 | 1,860 | 9,870 |
| *LMNB1* | 1 | 8,744 | 9,456 | 8,730 | 9,445 | 9,603 | 9,660 | 9,385 | 9,331 | 9,870 |

**Table S18.** Pareto-optimal knockdowns under the discovery signature, related to STAR Methods. These knockdowns show how multi-objective ranking behaves under the discovery signature. They are not proposed as rejuvenation targets. Sorted by bootstrap support (fraction of 200 donor bootstraps with the gene in the top 5%). Shifts are mean z over lines; n/m, target not measured; unknown, no annotation. After filtering (Figure S10B), the front is *ANLN*, *LSM12*, *GPS2*, *SMG7*, *ZRANB2*, *TFB1M*, *EIF4G1*, *SUMO2*, *SDHD*, *MRPS17*, *MRPS30*, *MTOR*, *HNRNPR*, *TERF1*, *SAE1*.

| **Gene** | **R<sub>rob</sub>** | **Sign consistency** | **Identity shift** | **Proliferation shift** | **Stress shift** | **Response size** | **On-target z** | **Cells (mean)** | **Bootstrap support** | **OncoKB tumor suppressor** | **CellAge** | **GenAge** | **Weak knockdown** |
|---|---|---|---|---|---|---|---|---|---|---|---|---|---|
| *ANLN* | 0.20 | 1.00 | 0.05 | −0.16 | 0.03 | 0.18 | −0.63 | 80 | 0.96 | no | inhibitor | unknown | no |
| *LSM12* | 0.16 | 1.00 | −0.05 | −0.06 | −0.01 | 0.11 | −0.91 | 81 | 0.78 | no | unknown | unknown | no |
| *UFL1* | 0.17 | 1.00 | 0.00 | −0.02 | 0.01 | 0.09 | −0.71 | 188 | 0.64 | no | unknown | unknown | no |
| *GPS2* | 0.16 | 1.00 | −0.04 | 0.01 | 0.00 | 0.09 | −0.76 | 142 | 0.46 | yes | unknown | unknown | no |
| *ZRANB2* | 0.14 | 1.00 | 0.02 | 0.01 | −0.01 | 0.12 | −1.21 | 61 | 0.42 | no | unknown | unknown | no |
| *ADNP2* | 0.14 | 1.00 | −0.01 | −0.08 | 0.01 | 0.15 | −0.48 | 49 | 0.33 | no | unknown | unknown | yes |
| *CRLS1* | 0.14 | 1.00 | 0.00 | −0.01 | 0.01 | 0.11 | −0.73 | 91 | 0.29 | no | unknown | unknown | no |
| *EGLN2* | 0.13 | 1.00 | 0.01 | 0.01 | 0.00 | 0.07 | −0.58 | 212 | 0.25 | no | unknown | anti-longevity | no |
| *TREML2* | 0.14 | 1.00 | −0.01 | 0.02 | 0.00 | 0.09 | n/m | 92 | 0.19 | no | unknown | unknown | yes |
| *CCDC144A* | 0.13 | 1.00 | 0.01 | 0.02 | 0.00 | 0.09 | n/m | 139 | 0.09 | no | unknown | unknown | yes |
| *OR4F4* | 0.13 | 1.00 | 0.01 | 0.00 | 0.00 | 0.12 | n/m | 64 | 0.09 | no | unknown | unknown | yes |
| *COQ5* | 0.12 | 1.00 | 0.00 | −0.02 | 0.00 | 0.11 | −0.52 | 127 | 0.06 | no | unknown | unknown | no |
| *MRPS25* | 0.05 | 0.75 | 0.00 | −0.02 | 0.05 | 0.11 | −1.05 | 101 | 0.00 | no | unknown | unknown | no |
| *NR2C2AP* | 0.04 | 0.75 | 0.00 | −0.11 | 0.03 | 0.15 | −0.38 | 74 | 0.00 | no | unknown | unknown | yes |
| *OR4K1* | 0.03 | 0.75 | 0.00 | 0.01 | 0.00 | 0.11 | n/m | 60 | 0.00 | no | unknown | unknown | yes |
| *NDUFB7* | 0.02 | 0.67 | 0.00 | −0.02 | 0.00 | 0.09 | −1.43 | 102 | 0.00 | no | unknown | unknown | no |

**Table S19.** Deviations from protocol v1.0 and later analyses, related to STAR Methods. Deviations (D) and later analyses (X), with dates. All X analyses were added after the main results were known. Output files for each row are listed in the repository.

| ID | Date | Change or analysis | Possible influence or status |
|---|---|---|---|
| Freeze | 12 October 2026 | Protocol v1.0 committed (commit 1eb8681, tag protocol-v1.0; SHA-256 01a573574e8225e5ba68ddd2757b81dcd765083274d9b3290b8a1bb0ca879fa7) | Unchanged since |
| D1 | 12 October 2026 | Orthologs from the Alliance of Genome Resources instead of Ensembl BioMart | Changes the mapped label set; not tuned to results |
| D2 | 12 October 2026 | The K562 essential-gene and genome-wide screens are averaged into one line for R<sub>rob</sub>, sign consistency and the Pareto ranking | Defines R<sub>rob</sub> |
| D3 | 12 October 2026 | H1 uses the K562 essential-gene screen as "K562"; H1b uses the 1,784 knockdowns measured in all screens rather than the 2,053 shared by the two K562 screens | None expected |
| D4 | 12 October 2026 | HepG2 and Jurkat pseudobulked by us; their non-targeting controls are 100 random groups of non-targeting cells | Normalization may differ slightly from the published release |
| D5 | 12 October 2026 | Ridge penalty λ = 1 for the H3 models | Untuned; may understate the combined model |
| D6 | 12 October 2026 | A file-size check (files under 1,000 bytes were skipped) silently dropped the yeast ortholog table in the first H4b run; in the corrected run the table passed the check and H4b was rerun. The check now raises an error instead of skipping | H4b AUROC 0.52 (Holm *P* = 0.18) before, 0.51 (Holm *P* = 0.27) after; conclusion unchanged |
| D7 | 12 October 2026 | Early exploratory additions: early sensitivity run on ages 18 to 80 (replaced by F1, ages 20 to 79), adjusted within-K562 agreement, calibrated line-transfer model, single-screen evidence tests, on-target annotation | Labeled exploratory |
| D8 | 26 October 2026 | In the cohort analyses of 13 October 2026 (X01 and X05), the GTEx-skin G0 model also contained ischemic time by mistake. G0 is now age + sex for both GTEx tissues; ischemic time enters the skin models from G1 | Skin G0 values changed (0.21 to 0.15; fibroblasts versus skin 0.02 to 0.06). No main result changed |
| D9 | 26 October 2026 | Exclusion flow written out: 143 GEO samples = 133 healthy donors + 10 progeria samples; 22 donors under 18 excluded, leaving 111 donors aged 19–96 years (GEO age field) | None |
| D10 | 26 October 2026 | Loader fix: the fibroblast loader read all sample columns except the last and so dropped one column (one donor aged 17). All sample columns are now read | None: the donor is under 18 and excluded by the age filter |
| D11 | 28 October 2026 | Identity genes: log-normalized gene mean of a line minus the mean of the other lines, among genes above that line's median expression, top 100 (protocol: z-score of the line's mean relative to the other lines on control expression). K562 and RPE1 use the gene means distributed with the Replogle release; HepG2 and Jurkat use non-targeting cells only | Defines the identity program ı used by V1, V3 and V7 |
| X01 | 13 October 2026 | GEO metadata (platform, repository, biopsy site); cohort sensitivity F0–F7 × G0–G2 with donor-bootstrap CIs and split-half references | Exploratory |
| X02 | 13 October 2026 | Score definitions V2–V5 | Exploratory |
| X03 | 13 October 2026 | Network baselines and CIs; calibrated line transfer | Exploratory |
| X04 | 13 October 2026 | Evidence coverage, label permutations, label subsets and proliferation-only baseline | Exploratory |
| X05 | 13 October 2026 | 2,000 donor bootstraps for every cohort specification (500 split halves) | Exploratory |
| X06 | 13 October 2026 | 2,000 donor bootstraps for H5 (the 200 bootstraps in the protocol give the main result) | Exploratory |
| X07 | 13 October 2026 | 500 degree-preserving randomized networks for H3a (the 10 nulls in the protocol give the main result); 2,000 bootstrap resamples for all CIs; 10,000 label permutations | Exploratory |
| X08 | 26 to 27 October 2026 | Sex as a positive control (all genes, autosomes, X/Y, top-200 genes); sign check of pre-named age genes and canonical sex genes | Exploratory |
| X09 | 26 to 27 October 2026 | Permuted-age null (1,000 permutations, main model); power-matched 74-donor subsets; age per repository; variance inflation factor of age | Exploratory |
| X10 | 26 to 27 October 2026 | F8 (without AG donors aged 80 or older), F9 and G3 (five hidden factors); 2,000 donor bootstraps for all specifications; bias-corrected intervals (27 October) | Exploratory |
| X11 | 26 to 27 October 2026 | 100 random expression-matched signatures through H1, H2 and H4 | Exploratory |
| X12 | 26 to 27 October 2026 | Split-cell reliability (HepG2, Jurkat; 20 splits); test–retest reliability (two K562 screens); size-matched non-targeting groups; disattenuated agreement; proliferation measured in held-out cells | Exploratory |
| X13 | 26 to 27 October 2026 | Paired DeLong tests, incremental logistic regression, minimum detectable AUROC; GenAge mouse-only and without yeast; label overlap with Hart CEGv2 and (27 October) DepMap 24Q2 common essentials | Exploratory |
| X14 | 26 to 27 October 2026 | R<sub>rob</sub> for single-line genes (equals R<sub>adj</sub>); H4 and H5 on genes measured in at least two lines | Exploratory |
| X15 | 26 to 27 October 2026 | In-fold network analysis: in-fold residualization, inner-CV ridge penalty, 10 CV seeds, common subset per screen; nearest-neighbor, gene-covariate, physical-network, calibrated-network and proliferation baselines; block bootstrap over response clusters | Exploratory |
| X16 | 26 to 27 October 2026 | Proliferation deciles, distance correlation, mutual information; definitions V6 (spline) and V7 (no magnitude); filtered Pareto front; ranks of senescence genes; up-minus-down score | Exploratory |
| X17 | 26 to 27 October 2026 | Cross-cohort elastic-net age prediction | Exploratory |
| X18 | 26 to 27 October 2026 | TRACE run end to end with a GTEx-derived discovery signature | Exploratory |
| X19 | 28 October 2026 | Permuted-age null for every discovery specification (F0–F9) against GTEx G0, both tissues; empirical two-sided P for genome-wide and top-200 agreement | Exploratory |
| X20 | 28 October 2026 | Permuted-sex null for the sex positive control; planted shared age-like effect (500 genes, two effect sizes, 200 replicates each) on age-permuted data | Exploratory |
| X21 | 28 October 2026 | Discovery hidden factors: association with age, repository and platform; Hallmark enrichment of the factor loadings | Exploratory |
| X22 | 28 October 2026 | Network baselines rerun with a wider ridge grid (10<sup>−6</sup> to 10<sup>3</sup>) and a nearest-neighbor predictor that uses the gene's response in the other lines only; the own-profile predictor kept as an upper reference | Exploratory |
| X23 | 28 October 2026 | GTEx-derived signature (X18): proliferation-only baseline, paired AUROC tests, oxidative-phosphorylation and stress program tests, Hallmark composition | Exploratory |
| X24 | 28 October 2026 | Random-signature null compared with the real signature without proliferation genes (V2 input) | Exploratory |
| X25 | 28 October 2026 | CellAge labels by manipulation method (CellAge web table); V6 and V7 effects on the top-5% lists and the Pareto front; table of all exploratory tests with family sizes | Exploratory |
| X26 | 29 October 2026 | Random-signature null: every raw-score statistic compared with the real signature without proliferation genes (V2 input), with one-sided empirical *P* | Exploratory |
| X27 | 29 October 2026 | Agreement among the k strongest discovery age genes, k = 50 to 1,000 (the choice k = 200 was made after the first results) | Exploratory |
| X28 | 29 October 2026 | Permuted-age null for the agreement between GTEx fibroblasts and GTEx skin | Exploratory |
| X29 | 29 October 2026 | Per-family Holm and Benjamini–Hochberg values for every exploratory test; report table with one row per check | Exploratory |
| X30 | 3 November 2026 | Permuted-age null (1,000 permutations) for agreement among the k strongest discovery genes at every k (50 to 1,000) and for the sign agreement of signature genes | Exploratory |
| X31 | 3 November 2026 | Hidden factors computed after a sex-only model (F9b), their correlation with age, and top-200 agreement with its own permuted-age null | Exploratory |
| X32 | 3 November 2026 | Planted shared effect added to the real (not permuted) data of both cohorts (planted part alone with an absolute t of about 4 and 6; 200 replicates each) | Exploratory |
| X33 | 3 November 2026 | Variance inflation factor of age in the GTEx batch models; Box 1 checks applied to this study (pass, fail or flag per check); duplicate rows removed from the exploratory-test table | Exploratory |
| X34 | 6 November 2026 | Permuted-age null for the GTEx batch model G2: age permuted within nucleic-acid isolation batch in GTEx and across donors in the discovery cohort (1,000 permutations), G2 refitted, top-200 agreement for F0 | Exploratory |
| X35 | 6 November 2026 | Box 1 cohort check for the GTEx-derived signature: the 200 strongest GTEx-fibroblast age genes against the discovery cohort, with the same 1,000 permuted age labelings as the F0 test | Exploratory |
| X36 | 6 November 2026 | Box 1 report for both signatures; replicate rule changed to reliability of the adjusted score in every screen and adjusted between-line agreement above the random-signature null; F9 t-statistics on the F0 top 200 | Exploratory |
| X37 | 6 November 2026 | Like-for-like cohort transfer: ranking rebuilt from discovery donors aged 20–79 (the GTEx age range) and compared with the GTEx-fibroblast ranking (no P value), with 100 donor bootstraps of the 20–79 subset as reference; Box 1 positive example: GTEx fibroblasts split into two random halves treated as two cohorts, with the cohort (200 permuted-age labelings), effect-size and stability (100 donor bootstraps) checks applied to split 0 and point estimates over 20 splits | Exploratory |
| X38 | 6 November 2026 | Box 1 report: label check for the GTEx-derived signature and the sex control use BH q within their Table S5 families instead of separately computed corrections, and the planted check is marked not run for the GTEx-derived signature (planted in the discovery direction only); results unchanged | Exploratory |

**Table S20.** Box 1 checks applied to this study, related to Box 1. Rules as in Box 1, applied to the discovery signature and to the GTEx-derived signature. Pass, the ranking can be trusted on this point; fail, it cannot without further evidence; flag, a warning that must be reported with the ranking; info, reported without a rule; not run, the input was not computed for that signature. For the GTEx-derived signature, the cohort check uses its own 200 strongest genes against the discovery cohort; the sex control, planted effect and power checks use the same cohorts, labels and pipeline as for the discovery signature. Its reliability is available only as K562 test–retest (one screen), so its replicate result rests on that screen; its label tests are exploratory and use BH q within their families in Table S5, including the paired DeLong comparison with the proliferation-only baseline. Corrected P, Holm-adjusted P for tests fixed before the study and Benjamini–Hochberg q for other tests. The stability rule and the AUROC-above-0.5 part of the label rule follow tests fixed before this study (Table 1); the comparison with the proliferation-only baseline and the other thresholds are suggestions set after this study and have not been tested on other screens. Produced by the TRACE command in Box 1.

| Item | Check | Threshold | Value, discovery | Result, discovery | Value, GTEx-derived | Result, GTEx-derived |
|---|---|---|---|---|---|---|
| Cohorts | Age effects, 200 strongest genes of the signature's cohort, versus the other fibroblast cohort (Spearman ρ; main model; for the discovery signature G1 and G2 for information) | above the permuted-age 97.5th percentile under the main model | −0.55 (G1 −0.46, G2 −0.17); null 97.5th percentile 0.43 | fail | −0.11; null 97.5th percentile 0.46 | fail |
| Cohorts | Planted shared effect added to the real data: median \|t\| of the planted part alone (against the median \|t\| of the observed top 200); runs with top-200 ρ above the null | planted \|t\| no larger than the observed median \|t\|, recovered in at least 0.80 of runs | \|t\| 4.0 (observed 6.2); 1.00 | pass | – | not run |
| Cohorts | Positive control: sex effects, 200 most sex-biased genes, versus GTEx fibroblasts (permuted-sex P; BH q within the sex-control family, Table S5) | corrected *P* < 0.05 | ρ = 0.65; *P* = 0.002; q = 0.012 | pass | same cohorts and pipeline | pass |
| Cohorts | Positive control: sex effects, 200 most sex-biased genes, versus GTEx skin (permuted-sex P; BH q within the sex-control family, Table S5) | corrected *P* < 0.05 | ρ = 0.32; *P* = 0.057; q = 0.17 | fail | same cohorts and pipeline | fail |
| Programs | Raw score versus proliferation shift (Spearman ρ, range over screens); real signature versus 100 random signatures (discovery only) | flag if \|ρ\| ≥ 0.3 in any screen or the real signature exceeds 95% of random signatures | 0.49 to 0.90; above all random signatures in every screen | flag | −0.39 to 0.07 | flag |
| Replicates | Raw score: within-line (two K562 screens) and mean between-line agreement, side by side (one-sided 95% bound of the difference) | reported | 0.50 and 0.51 (−0.04) | info | 0.45 and 0.33 (0.08) | info |
| Replicates | Adjusted score (V1): reliability (range over screens; split cells in HepG2 and Jurkat, test–retest in K562) | ≥ 0.5 in every screen | 0.38 to 0.64 | fail | K562 test–retest 0.44 (one screen only); split cells not run | fail |
| Replicates | Adjusted score: median between-line agreement against 100 random signatures (real signature without proliferation genes; one-sided empirical P) | above the random-signature null (*P* < 0.05) | 0.21 (null maximum 0.18; *P* = 0.010) | pass | – | not run |
| Labels | CellAge: R<sub>rob</sub> (V1) AUROC, all genes, and proliferation shift alone (discovery: Holm P, fixed in advance; GTEx-derived: BH q, exploratory, with paired DeLong q) | AUROC above 0.5 (corrected *P* < 0.05) and above the proliferation-only AUROC (paired DeLong, corrected *P* < 0.05) | 0.54 (*P* = 0.10); baseline 0.58 | fail | 0.58 (q = 0.0092); baseline 0.58 (paired q = 0.94) | fail |
| Labels | GenAge: R<sub>rob</sub> (V1) AUROC, all genes, and proliferation shift alone (discovery: Holm P, fixed in advance; GTEx-derived: BH q, exploratory, with paired DeLong q) | AUROC above 0.5 (corrected *P* < 0.05) and above the proliferation-only AUROC (paired DeLong, corrected *P* < 0.05) | 0.51 (*P* = 0.27); baseline 0.42 | fail | 0.53 (q = 0.27); baseline 0.42 (paired q = 0.0013) | fail |
| Power | Minimum detectable AUROC at 80% power, CellAge | a label test above 0.60 is reported as uninformative | 0.56 | info | 0.56 | info |
| Power | Minimum detectable AUROC at 80% power, GenAge | a label test above 0.60 is reported as uninformative | 0.55 | info | 0.55 | info |
| Power | Minimum detectable AUROC at 80% power, GenAge, mouse genes only | a label test above 0.60 is reported as uninformative | 0.67 | uninformative | 0.67 | uninformative |
| Effect sizes | Top-5% overlap between the ranking and the ranking from the other fibroblast cohort | at least half the overlap between donor bootstraps of one cohort | 0.04 (bootstraps 0.47) | fail | 0.04 (bootstraps 0.40) | fail |
| Effect sizes | Top-5% overlap between the ranking and the ranking from GTEx skin | at least half the overlap between donor bootstraps of one cohort | 0.11 | fail | 0.04 | fail |
| Knockdown quality | Pareto-front genes with weak or unmeasured on-target knockdown before filtering; front after filtering | report the share, then rank only after filtering | 6 of 16; 15 genes after filtering | info | 16 of 47; filtering not run | info |
| Stability | Median donor-bootstrap ρ of the ranking (all genes; genes in ≥ 2 lines) | ≥ 0.8 | 0.779; 0.856 | fail (all genes); pass (genes in ≥ 2 lines) | 0.671; not run | fail (all genes) |

